# Learning Binding Affinities via Fine-tuning of Protein and Ligand Language Models

**DOI:** 10.1101/2024.11.01.621495

**Authors:** Rohan Gorantla, Aryo Pradipta Gema, Ian Xi Yang, Álvaro Serrano-Morrás, Benjamin Suutari, Jordi Juárez-Jiménez, Antonia S. J. S. Mey

## Abstract

Accurate *in-silico* prediction of protein-ligand binding affinity is essential for efficient hit identification in large molecular libraries. Commonly used structure-based methods such as docking often fail to rank compounds effectively, and free energy-based approaches, while accurate, are too computationally intensive for large-scale screening. Existing deep learning models struggle to generalize to new targets or drugs, and current evaluation methods often do not accurately reflect real-world performance. We introduce **BALM**, a deep learning framework that predicts **b**inding **a**ffinity using pre-trained protein and ligand **l**anguage **m**odels. We also propose improved evaluation strategies with diverse data sets and metrics to assess model performance to new targets better. Using the BindingDB dataset, BALM generalises unseen drugs, scaffolds, and targets. In few-shot scenarios for targets such as *USP7* and *Mpro*, it outperforms traditional machine learning and docking methods, including AutoDock Vina. Adoption of our target-based evaluation methods will allow a more stringent evaluation of machine learning-based scoring tools. Our protein prediction framework shows good performance, is computationally efficient, and is highly adaptable within this evaluation setting, making it practical for early-stage drug discovery screening.

## Introduction

Identifying hit compounds from ultra-large libraries for accelerating target-based drug discovery hinges on the precision of *in-silico* methods for predicting protein-ligand binding affinity (BA). Structure-based approaches such as giga-docking^1,2^ are commonly used for virtual screening.^3,4^ While docking is often a good approach to generate chemically reasonable poses, it usually falls short in rank-ordering compounds.^5^ Free energy-based methods^6–8^ offer more reliable predictions but are computationally costly for hit discovery in ultra-large libraries and are usually restricted to lead optimization stages. Limited size and diversity of screening libraries have long been a bottleneck for the detection of novel potent ligands and for the whole process of drug discovery. ^9^ With recent advancements in on-demand libraries such as Enamine, ^4,10^ scaling up the virtual screening capabilities from million to a trillion compounds is possible. The chances of these libraries containing more potent ligands with better physicochemical properties increase by increasing the search-space. However, the number of potential decoys that need to be filtered out will also increase. Effective hit identification in ultra-large libraries requires computationally efficient and accurate *in-silico* methods. Machine learning advances, particularly deep learning (DL), accelerate drug discovery stages, from virtual screening^11–15^ to finding hits to optimizing lead compounds with active learning^16–18^ and *de-novo* design^19–22^ techniques.

In recent years, many DL strategies have been developed for predicting binding affinity,^12,13,23–26^ yet challenges remain at the model, data, and evaluation levels. At the model level, DL methods typically fall into two categories - complex-based and sequence-based. Complex-based models leverage three-dimensional (3D) protein-ligand structural data, often derived from databases such as PDBBind.^27^ Although the use of 3D structures can potentially capture detailed interaction information, complex-based models often fail to generalize due to the limited availability of high-quality 3D complexes and biases within structural data.^28^ On the sequence-based side, models are trained using 1D protein sequences and SMILES representations of ligands. This makes them more flexible in terms of data requirements due to large sequence databases but is limited to a much smaller pool of interaction data in comparison to typical language model training datasets. Furthermore, investigations into understanding what these models learn from input protein sequence and ligand SMILES data have shown a dependence on ligand information and neglect of combined information on protein and ligand for prediction.^29^ Challenges at the data and evaluation levels further hinder DL model performance in binding affinity prediction. Commonly used datasets for training these DL models combine IC50 and *K*_i_ data from various assays, introducing significant noise due to differences in how experiments are performed.^30^ This undermines the reliability of the model. Furthermore, evaluating these models by randomly splitting data across train and test sets often results in data leakage, where similar compounds end up in both training and test sets, leading to overestimating the model’s generalizability.^31^ We address these three key challenges at **model, data,** and **evaluation levels** to provide a DL framework for practical screening purposes.

At the model level, we introduce **BALM**, a novel sequence-based deep learning method for predicting protein-ligand **b**inding **a**ffinity using pre-trained protein and ligand **l**anguage **m**odels. The key novelty over previous metric learning-based methods for binding affinity prediction,^36–38^ is that BALM learns binding affinities based on the distance between proteins and ligands in feature space. BALM operates on pairs of data (protein sequences, ligand SMILES) as shown in Fig. 1 to learn an embedding space that directly represents the binding affinity pK*_d_*. BALM builds on advances in protein^32^ and ligand^33^ language models, which learn representations in an unsupervised manner from large-scale datasets such as UniRef^39^ for proteins and PubChem^40^ for ligands. BALM uses these protein (ESM-2) and ligand (ChemBERTa-2) language models to extract features and incorporates parameter-efficient fine-tuning (PEFT)^41–46^ to adapt these embeddings specifically for binding affinity prediction. This is achieved by tuning only a small fraction of model parameters and significantly reducing computational costs for training. By embedding protein-ligand pairs in a shared feature space that reflects interaction strength, BALM optimizes the cosine similarity (a distance metric) to directly predict binding affinity (*pK*_d_).

**Figure 1:**
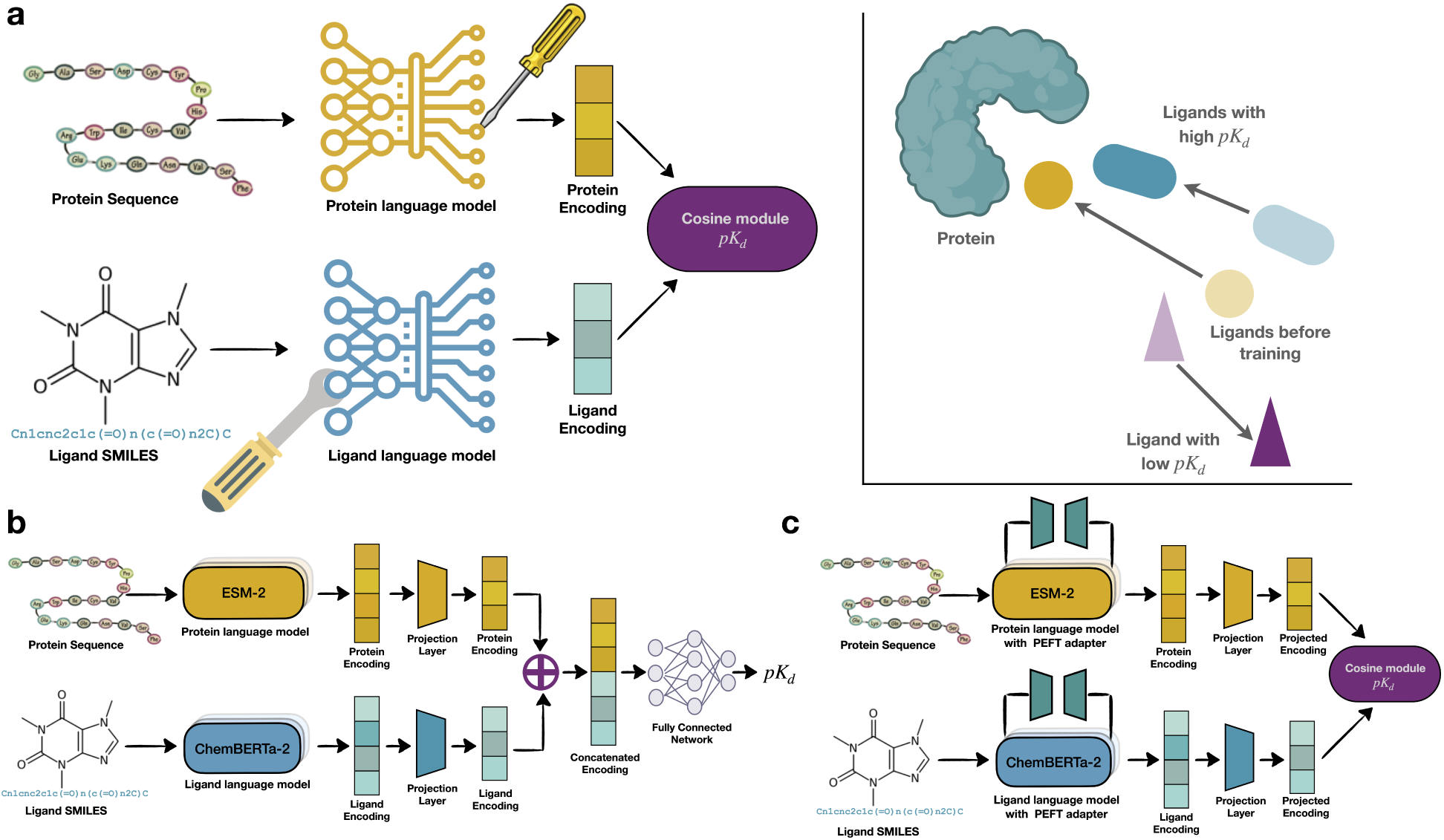
Overview of BALM architecture. **(a)** BALM learns by optimizing the distance between protein and ligand encodings using a cosine similarity metric directly representing binding affinity (p*K*_d_). Cosine similarity is maximized for binding interactions and minimized for non-binding interactions. Protein sequences and ligand SMILES strings are encoded using language models trained on extensive protein and ligand databases. ESM-2^32^ is used for encoding protein sequences, and ChemBERTa-2^33^ for extracting features from ligand SMILES. These encoded features are projected into a shared latent space via linear layers. The model then optimizes the cosine similarity between protein and ligand embeddings to learn binding affinity. **(b)** The baseline model is based on a pipeline from previous studies ^13,18,24,34,35^ where features are first extracted from protein sequences and SMILES strings using a deep learning architecture. Then these features are concatenated and passed through a linear layer to predict binding affinity. The model is trained using a Mean Squared Error (MSE) loss function to predict the binding affinity. **(c)** BALM with parameter-efficient fine-tuning (PEFT). PEFT adapters are added to the protein (LoKr) and ligand (LoHa) language models, allowing selective fine-tuning of small subsets of additional parameters while keeping the main model weights frozen. The fine-tuned embeddings are then projected and optimized using cosine similarity similar to the original BALM for affinity prediction.

To ensure the best data integrity from the BindingDB dataset, we only use *K*_d_ values for training and remove any assay limits in the standard dataset. While reducing the size of the training set, this makes it of higher quality. In terms of improved evaluation strategies, we systematically test BALM’s performance across challenging data splits within the BindingDB dataset,^47,48^ including zero-shot (cold drug and cold target), scaffold, and random splits. Our experiments demonstrate that BALM outperforms our literature-inspired baseline model across all splits, showcasing generalization to unseen drugs, scaffolds, and targets on typically reported aggregate evaluation metrics such as RMSE and correlation coefficients. Incorporating PEFT further enhances BALM’s performance, highlighting the benefits of parameter-efficient fine-tuning for producing reliable affinity models.

Crucially, the true picture of a model’s utility in a drug discovery scenario cannot be evaluated by reporting its performance on aggregated protein targets.^8^ We evaluate zero-shot predictions across individual protein targets and suggest using Fisher-transformed correlations to capture target-specific performance. This metric captures the significant variability in zero-shot predictions across individual protein targets, particularly in cold target settings where protein-ligand data are sparse. By utilizing Fisher-transformed^49^ correlations to capture target-specific performance, we obtain a clearer view of BALM’s practical applicability and limitations, offering a model evaluation approach that could inform the development and fair assessment of future DL methods. To further assess BALM’s adaptability, we evaluated its performance in practical few-shot learning scenarios on two unknown (to the model) targets—*USP7*, a ubiquitin-specific protease,^50^ and *Mpro*, the main protease of SARS-CoV-2.^51,52^ Using a pre-trained BALM model, we fine-tuned the embeddings for these targets, observing rapid and notable performance gains with only a small fraction of additional data, highlighting BALM’s potential in real-world settings with limited experimental data. We benchmarked BALM against simple machine learning models, such as Gaussian Processes, which depend heavily on feature selection and kernel choice, impacting generalizability and performance. BALM’s performance is robust across different targets without further tuning embedding requirements, emphasizing its practicality in real-world screening scenarios. We further compared BALM’s performance with traditional structure-based docking methods, including AutoDock Vina^53^ and rDock,^54^ on the Leak Proof PDBBind dataset,^55^ which was designed to eliminate test set data leakage. BALM outperformed these methods in the rescoring of crystal structures in predictive accuracy. Lastly, we evaluated pre-trained BALM on the Free Energy Benchmark,^8^ which includes data representative of the lead optimization stage, featuring congeneric series for three challenging targets (*MCL1*, *HIF2A*, *SYK*). While BALM struggled on *HIF2A* and *SYK*, it showed competitive accuracy by few-shot fine-tuning on the larger *MCL1* dataset. For highly similar ligands, as in lead optimization datasets, BALM’s predictions cluster within a narrower range as already observed in other models,^26^ pointing towards some current limitations. This work demonstrates how, under rigorous evaluation metrics, large language models represent an attractive solution for fast and accurate protein property prediction.

## Results

### BALM learns better than baseline models across challenging data splits

We investigated the performance of the BALM model (Fig. 1a) on the BindingDB dataset^47,48^ and compared it with a consensus Baseline model^13,18,24,34,35^ using different data splits. We focused on the BindingDB dataset, specifically looking at binding affinities measured according to *K*_d_,^48^ which contains around 48,000 non-covalent interactions involving 1,090 protein targets and 9,900 ligands. To ensure stable model training, we transformed the K_d_ values into *pK*_d_ values, following recommendations from previous studies. ^24,29,35^ To address significant bias caused by binding assay limits and the resulting skewed range of affinity predictions (Fig. S1), we cleaned the dataset by removing assay limits. This resulted in discarding around 21,000 interactions around a *pK_mathrmd_* of 5. After this cleaning step, the dataset was reduced to about 25,000 interactions, involving around 1,070 targets and 9,200 ligands. To evaluate the efficacy of each model, we considered random, cold target, cold drug, and scaffold dataset splits, as discussed in more detail in the data section. For the random split, interactions were randomly divided into train and test sets. The cold target split involved grouping protein targets into distinct sets to assess performance on unseen proteins, while the cold drug split assigned ligands similarly to test novel compounds. For the scaffold split, drugs were categorized by their two-dimensional Murcko scaffolds^56^ using RDKit’s `MurckoScaffold` module,^57^ testing the model’s ability to generalize to new chemical scaffolds. Target and drug-specific data splits are key for generalizability. We evaluated the models’ ability to generalize using the concordance index (CI), Pearson correlation, Spearman rank correlation, and root mean squared error (RMSE). To compare the performance of two trained models on the test set, we assessed statistical significance using paired t-tests, with *p <* 0.05 considered significant.

The Baseline model (Fig. 1b) follows a pipeline from previous studies ^13,18,24,34,35^ where features are extracted from protein sequences and SMILES strings using a DL architecture. These features are then concatenated and passed through a linear layer to predict binding affinity, with the Mean Squared Error (MSE) loss function used to learn *pK*_d_. For a fair comparison, we employed protein and ligand language models and utilized projected encodings from the projection layer similar to BALM. These encoded representations are concatenated and passed through a fully connected layer to predict binding affinity. Both the baseline and BALM models were run with three random seeds to ensure the model performance was consistent with random variation in the data splits or initialization.

Fig. 2a provides an overview of the evaluation metrics performance of the BALM (medium blue) and Baseline (light blue) models across four data splits. In the Random split, BALM outperforms the baseline model in all evaluation metrics with statistical significance.

**Figure 2:**
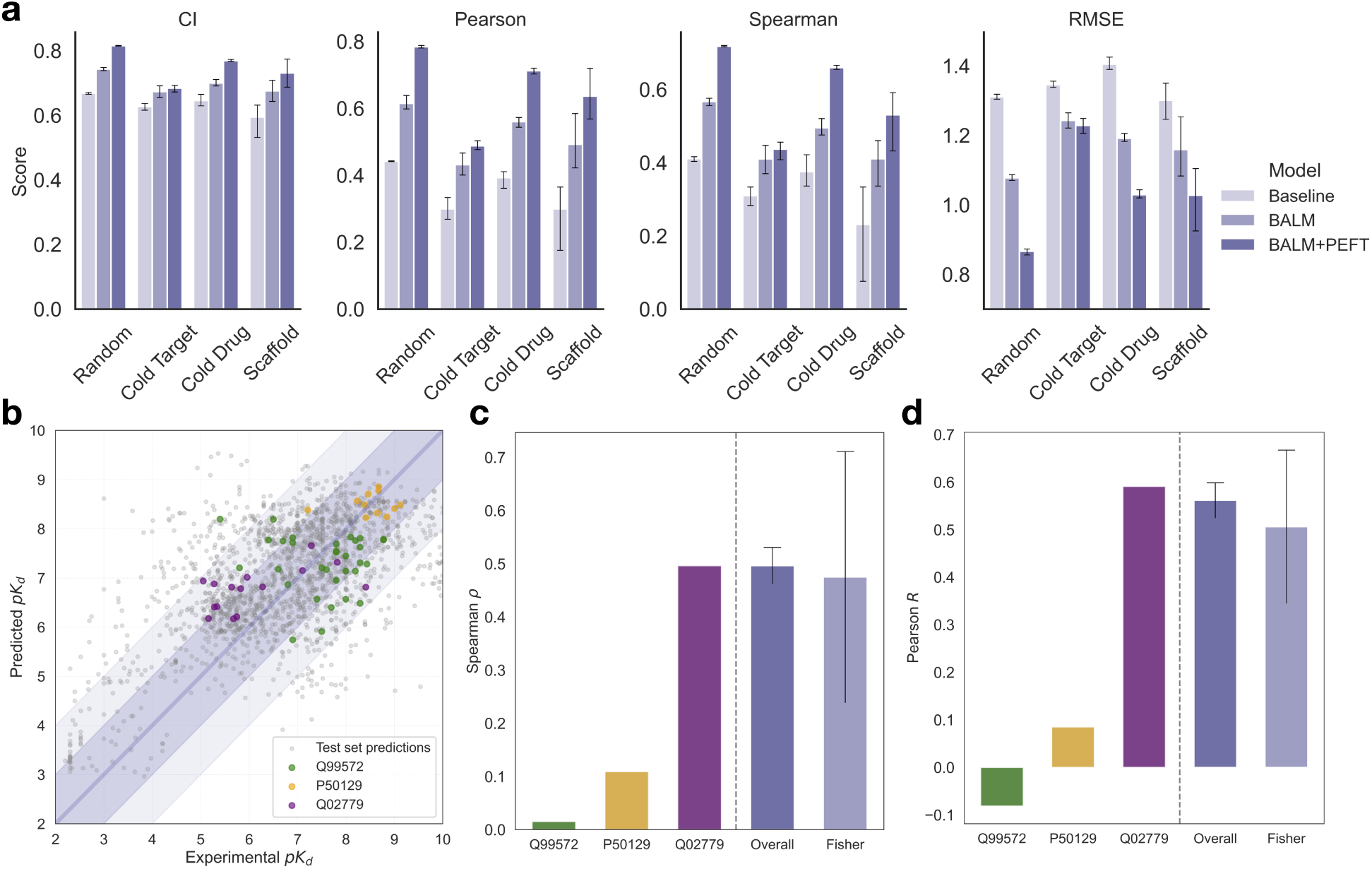
BALM performs better than the baseline model on challenging data splits, and parameter-efficient fine-tuning further improves its performance. **(a)** Performance comparison of Baseline, BALM, and BALM+PEFT models on the filtered BindingDB dataset using four data splits - Random, Cold Target, Cold Drug, and Scaffold. Error bars indicate the standard deviation that was observed during three random variations in the data splits and model initializations. Across all splits, BALM+PEFT (dark blue) demonstrates the best performance, as reflected by increased Pearson correlation and reduced RMSE. **(b)** Scatter plot showing zero-shot predictions of the BALM+PEFT variant on three randomly selected targets from the test set in cold target split. Grey points represent predictions across the entire test set, while coloured points highlight specific targets: *Q99572* (green), *P50129* (yellow), and *Q02779* (purple). **(c)** Spearman correlation (*ρ*) bar plot for selected test targets compared to overall dataset performance and Fisher transformed mean performance. *Overall ρ* indicates the model’s correlation calculated across all test set predictions, while the *Fisher ρ* aggregates individual correlations for each target by averaging correlations at the target level. The *Fisher ρ* provides a more realistic picture of model performance, accounting for target-specific variability and showing higher variance, as some targets may have better predictions (high Spearman) while others show worse performance (low Spearman). Larger standard deviations in the *Fisher transformed* values reflect these differences across targets. **(d)** Pearson correlation (*R*) bar plot for selected test targets, similar to panel (c). The *Overall R* reflects the model’s global performance across the test set, while the *Fisher* transformed aggregated *R* gives a view of target-specific correlations. *Fisher* aggregation helps highlight the performance distribution across targets, and large standard deviations for the *Fisher* suggest that some targets are predicted well while others are predicted poorly, indicating variability in the model’s zero-shot performance across unseen targets.

While the performance of BALM was in general better than the Baseline across all splits and metrics, the results on the most challenging splits were underwhelming.

Previous studies ^44–46^ have demonstrated that parameter-efficient fine-tuning (PEFT) can tailor the performance of pre-trained language models for systems not used during training by updating a small fraction of the parameters. We investigated the use of four different parameter-efficient adapters (LoRA, LoHa, LoKr, and IA3), which are added to the language models as shown in Fig. 1c., to further improve the performance of BALM. Using the BindingDB data with random splitting, we evaluated separately the effect of the different PEFT adapters on the ligand and the protein language models (Fig. S2), using three different values of the rank matrix hyperparameter *r* (8, 16 and 32) for the three methods that required this hyperparameter. LoHa (*r*= 16) showed the highest performance boost for ligand-only fine-tuning, improving Pearson correlation by 9.4%, while LoKr (*r*= 8) protein-only fine-tuning increased the value of the Pearson correlation by 18.2%. Combining the best-performing methods for both ligand and protein yielded the BALM+PEFT model, which demonstrated a substantial 23.4% improvement over the performance of the non-optimized version of BALM. Beyond the random split, the BALM+PEFT variant consistently outperformed the initial model across all splits and metrics (Fig. 2a). Improved performance in the cold target and cold drug splits are noteworthy, with increases in Pearson correlation of approximately 20% and 10% respectively and a decrease of ca. 5% in RMSE in both cases.

BALM+PEFT is a fine-tuned model that, according to the monitored metrics, displays high predictive power, and reliability, and seems highly generalisable to new targets and chemical scaffolds, as shown by its performance in the cold protein and cold target splits. However, evaluating the global performance of these models, grouping different protein targets and mixing extremely different chemical scaffolds does not truly inform the models’ performance in situations usually faced during drug discovery efforts. This is the ability to accurately screen millions of compounds of unknown affinity to a single, oftentimes previously unseen, macromolecular target. To this end, we investigated the performance of BALM+PEFT in the affinity prediction in the cold-target setting in more detail.

### Zero-shot performance is not reliable for all targets, and commonly used cumulative metrics do not give the real picture in cold target-setting

We evaluated the zero-shot performance of the BALM+PEFT model in the cold target split using selected targets to understand its performance on unseen proteins. Fig. 2b shows the zero-shot predictions for three targets of pharmacological relevance: the P2X purinoreceptor 7 (P2RX7, Uniprot ID: *Q99572*, the serotonin receptor 2A (HTR2A *P50129*), and the Mitogen-activated Protein kinase kinase kinase 10 (MAP3K10, *Q02779*). Notably, there is significant variability in the values of both Spearman and Pearson correlation coefficients with respect to experimental values for these protein targets, with values ranging close to 0 for the P2RX7 (green dots) and HTR2A (yellow dots) and very close to the average performance of the model in the MAP3K10 case (purple dots). Interestingly, employing the often-used cumulative correlations over the whole dataset seems to overestimate the reliability of the model in cold target-setting and fails to highlight target-dependent deviations, making them a potentially misleading metric of the overall model performance. In contrast, averaging Fisher-transformed correlation coefficients and then back-transforming the average to the Pearson coefficient can help identify the greater dispersion across different targets (see Supplementary information for more details). The variability in the performance of zero-shot predictions is also replicated outside the BindingDB data set, as demonstrated for the test cases of the Ubiquitin carboxyl-terminal hydrolase 7^50^ (USP7, *Q93009*) and the Main protease domain of the SARS-CoV-2 virus replicase polyprotein 1a (Mpro, *P0DTC1*).^51,52^ In both cases we transformed the available experimental IC50 into pIC50. For evaluation, we computed a change in free energy Δ*G* of binding for a final comparison between experimental and computed values. Since in the single target sencario experiments come from a single assay we can use the Cheng-Prusoff equation to convert to Δ*G*, please see the supplementary information for more details. For the two single targets we obtained significantly different performance of the BALM-PEFT model: zero-shot *r* = 0.64 and RMSE of 1.49 kcal/mol for the USP7 case (Fig. 3b) and *r* = 0.11, and the RMSE is higher at 2.11 kcal/mol for the Mpro (Fig. 3e), indicating that zero-shot performance inconsistencies are related to the model and not the dataset composition. One thing to note is that the homodimer stoichiometry is not taken into account in training or for the prediction of this model.

**Figure 3:**
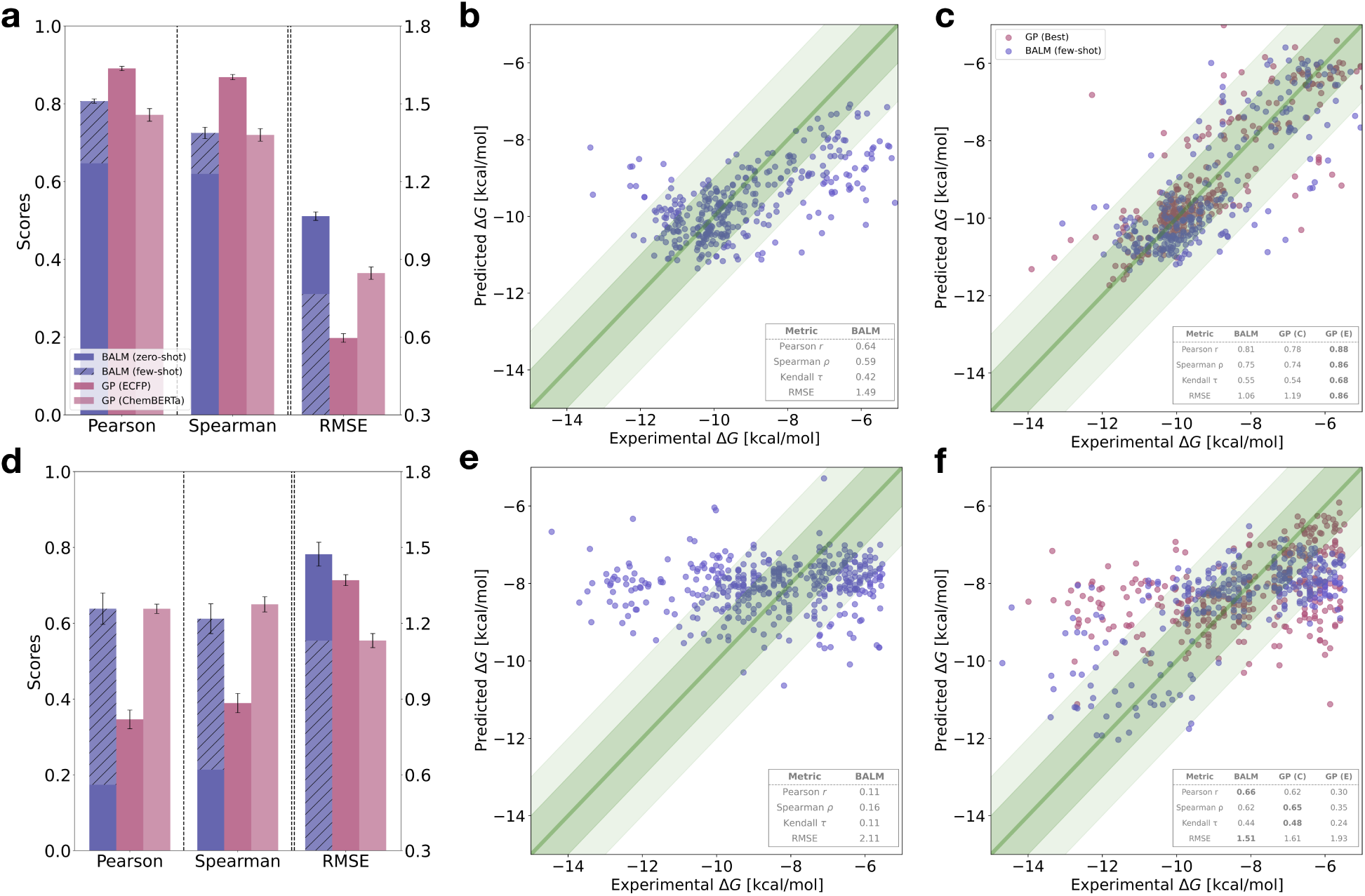
Zero-shot performance of pre-trained BALM+PEFT model and few-shot comparison of BALM with Gaussian Process (GP) models on *USP7* and *Mpro* targets. **(a, d)** Performance metrics for Pearson correlation, Spearman correlation, and RMSE on *USP7* (a) and *Mpro* (d) targets, comparing BALM in zero-shot (solid) and few-shot (patterned) settings with GP models trained using ECFP8 fingerprints (GP (E), Tanimoto kernel) and Chem-BERTa embeddings (GP (C), RBF kernel). The pre-trained BALM+PEFT model is fine-tuned by retraining only the projection layer. Error bars indicate standard deviation over three random seeds. **(b, e)** Scatter plots showing zero-shot model predictions of pre-trained BALM+PEFT model for *USP7* (b) and *Mpro* (e) targets. Experimental Δ*G* values (kcal/mol) are on the x-axis and predicted Δ*G* values are on the y-axis. Only 20% of the test set (selected randomly) is shown for readability. **(c, f)** Few-shot BALM+PEFT predictions (blue) and GP predictions (GP (Best), red) for *USP7* (c) and *Mpro* (f) targets, highlighting BALM’s robust performance across different GP configurations. We show 20% of the test set (selected randomly) for readability. GP (Best) refers to the optimal GP configuration for each target: GP (C) for *USP7* and GP (E) for *Mpro*, with metrics displayed for direct comparison.

### With few-shot fine-tuning, the BALM framework rapidly adapts to new targets and can be used for screening compound libraries

While zero-shot performance is highly variable depending on the target, it is common in current drug discovery efforts to have access to a small set of high-quality data points from the very early stages of the hit-to-lead process. We hypothesized that in a real-life scenario, these data points could be incorporated into a pre-trained model to improve its adaptability to targets beyond the training set. To explore this extent, we used the BALM+PEFT model trained on BindingDB as the base model and fine-tuned only the linear projection layer, which accounts for 262,656 out of 152.5 million total parameters, or approximately 0.17% of the model’s parameters. This should improve the adaptability of the model while being computationally efficient to implement. We compared the few-shot performance of BALM+PEFT with that of Gaussian Process (GP) models,^58^ which are surrogate models that can produce reliable predictions with limited data ^17,18^ only reling on ligand information. We used two molecular representations to train the GP models: Extended-Connectivity Fingerprints (ECFP8) paired with a Tanimoto kernel and ChemBERTa embeddings paired with a radial basis function (RBF) kernel.

We evaluated the few-shot performance on the *USP7* (Fig. 3a,b,c) and *Mpro* (Fig. 3d,e,f), datasets. On the one hand, BALM+PEFT exhibits notable improvements in Pearson and Spearman correlations and a reduction in RMSE with few-shot fine-tuning using 20% of the data for training and the rest for testing. All the models are trained and tested with three different random seeds as before to estimate variability from training. In the zero-shot setting, BALM’s Pearson correlation is 0.64 for *USP7* (Fig. 3a) and 0.11 for *Mpro* (Fig. 3d) that improves when trained on a few labelled examples to a Pearson correlation of 0.81 for *USP7* and 0.66 for *Mpro*, with RMSE values decreasing from 1.49 kcal/mol to 1.03 kcal/mol for *USP7* and from 2.11 kcal/mol to 1.51 kcal/mol for *Mpro*. On the other hand, the GP models, while displaying similar or even better accuracy than BALM+PEFT, are very sensitive to the choice of molecular representation, as shown in Fig. 3c and f. Specifically, in both datasets the Pearson correlation and RMSE worsen when switching from the ECFP8 fingerprint to the CheBERTa embedding. ECFP8+GP can be a naive starting point but will display more performance variability than a fine-tuned BALM+PEFT model. In contrast, the results obtained for BALM+PEFT demonstrate that, in a few-shot scenario, is a robust predictor independently of the choice of molecular representation. Furthermore, the model is very computationally efficient, with zero-shot predictions taking approximately 90 seconds per target for the 2,000 ligands in both USP7 and Mpro datasets on a single Nvidia A100 GPU. The few-shot approach required between 14 and 25 additional minutes for retraining using between 10% and 30% of the data. Refer to Fig. S3 for a detailed performance comparison of the pre-trained BALM+PEFT model (zero-shot) and few-shot fine-tuning using 10%, 20%, and 30% of experimental data on *USP7* and *Mpro* targets. As such, BALM+PEFT’s combination of zero-shot capability and robust performance with minimal additional data and fine-tuning on a small subset of parameters makes the model a robust tool for drug discovery screening scenarios, where rapid adaptation to novel targets and minimal dependency on feature engineering is essential to achieve the high-throughput required in current pharmaceutical industry settings.

### BALM achieves better ranking than docking scores for a variety of target families

To further establish the potential of BALM+PEFT-like models to be incorporated in virtual screening pipelines, we investigated the performance of the BALM+PEFT on the LP-PDBBind dataset^55^ and compared it to two well-established docking methods: AutoDock Vina^53^ and rDock.^54^ The LP-PDBBind dataset, which is derived from PDBBind v2020,^59^ consists of 2,100 protein-ligand complexes from 12 protein families with experimentally measured binding affinities. LP-PDBBind is specifically designed to minimize similarities in sequence, structure, and ligand chemistry across training, validation, and test sets, minimizing data leakage and allowing the testing of the model’s generalizability. For our evaluation, we employed the Clean Level 2 (CL2) split, as suggested by Li et al.,^55^ due to its higher structural reliability.

In Fig. 4, we compare the general performance of zero-shot BALM+PEFT with that of both docking methods. For both docking methods existing crystal structure poses were scored with the most appropriate scoring function (see methods). Results are presented as bar plots for Fisher-transformed Pearson (Fig. 4a) and Spearman correlations (Fig. 4b), as well as RMSE (Fig. 4c) across target families. BALM+PEFT consistently achieves higher Fisher-transformed Pearson and Spearman correlations compared to both docking methods, indicating stronger predictive and ranking performance across families. We also examined individual target families, comparing the performance of BALM+PEFT (in blue) versus Autodock Vina (Vina) (in orange) on three representative target families as identified in the LP-PDBBind dataset (Fig. 4d-f). For *Transferase* and *Chaperone* proteins (Fig. 4d), BALM+PEFT achieves Pearson correlation coefficients of 0.65 and 0.72 and RMSE values of 1.82 kcal/mol and 1,75 kcal/mol respectively, substantially outperforming Vina (Pearson r 0.12 / 0.65 and RMSE 6.22 kcal/mol / 1.94 kcal/mol). However, the ranking metrics for BALM (*ρ* = 0.58, *τ* = 0.42) and Vina (*ρ* = 0.57, *τ* = 0.41) are similar in the case of *Chaperone* proteins. For *Oxidoreductase* targets (Fig. 4f), Vina surpasses BALM+PEFT in ranking metrics (Spearman *ρ* = 0.66 vs. 0.59 for BALM), but BALM+PEFT achieves a lower RMSE of 1.73 kcal/mol versus Vina’s 2.88 kcal/mol. In the case of more challenging protein families, such as those containing metals, membrane proteins, or transcription factors both methods achieve comparable results, although in general RMSE values are lower for the BALM+PEFT predictions. A full comparison across the 12 families is provided in the supplementary information both for BALM+PEFT vs. Vina (Fig. S4) and BALM+PEFT vs rDock (Fig. S5). The results suggest that BALM+PEFT offers a fast and high-accuracy alternative to scoring functions and, as such, can be envisaged as a tool in virtual screening protocols as a way to re-rank docking solutions before progressing to more computationally demanding methods, such as alchemical free energy methods. Furthermore, the performance of docking methods is very dependent on the availability of high-quality three-dimensional protein-ligand complex structures, while BALM+PEFT bypasses the need for that, making it particularly appealing for targets or compounds for which is not possible to obtain reliable three-dimensional structures.

**Figure 4:**
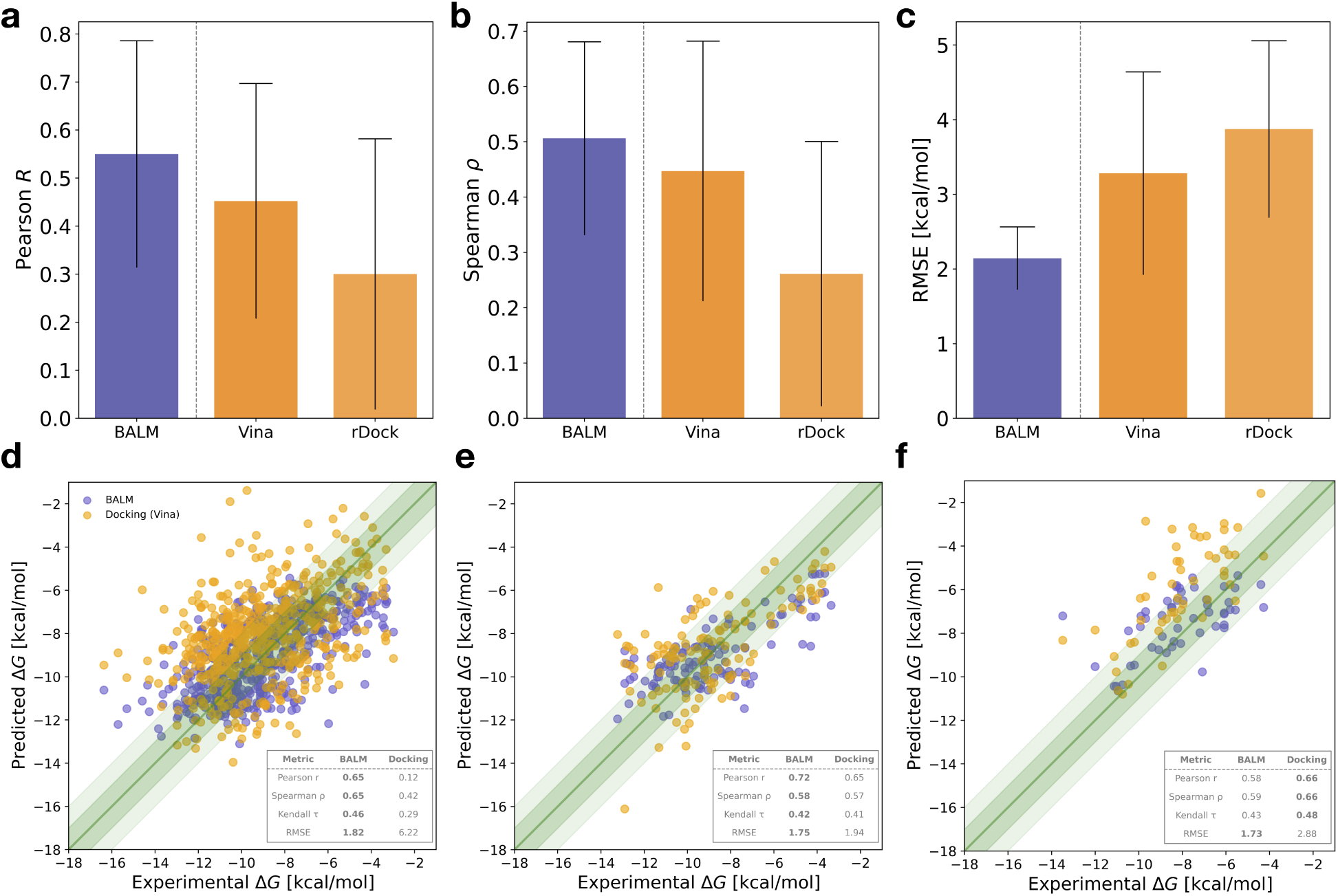
Evaluation of BALM with respect to docking methods, AutoDock Vina ^53^ and rDock, ^54^ across various target families in the LP-PDBBind^55^ test split. The test set includes approximately 2,100 protein-ligand complexes spanning 12 diverse target families, with results shown here for representative families. Fisher-transformed metrics are used for comparing Pearson and Spearman correlations across target families, providing an accurate performance measure that accounts for variability within each family. AutoDock Vina consistently performs better than rDock across most target families. **(a)** Bar plot of Pearson correlation for each method across all families; error bars represent the standard deviation across target families. **(b)** Spearman correlation, showing the ranking accuracy of each method across families with similar error bars. **(c)** RMSE, directly comparing the predictive accuracy of each method, where lower values indicate higher precision in binding affinity predictions. **(d-f)** Scatter plots showing the performance of BALM (blue) versus AutoDock Vina (orange) for different target families. **(d)** For *Transferase* targets (545 PDB complexes), BALM achieves superior Pearson and Spearman correlations compared to Vina, demonstrating its effectiveness in predicting kinase-related binding affinities. **(e)** For *Chaperone* targets (119 PDB complexes), BALM outperforms Vina, especially in RMSE, which suggests higher prediction stability for this family. **(f)** In the case of *Oxidoreductase* targets (52 PDB complexes), docking methods show slightly better ranking performance, while BALM achieves a lower RMSE, indicating more precise binding affinity predictions for this target class.

### BALM on free energy benchmark datasets

Last, we compared the performance of BALM+PEFT with alchemical free energy (AFE) methods for the evaluation of relative binding free energies.^7^ We used a subset of targets from the protein-ligand free energy benchmark curated by Hahn et al.,^8^ which is composed of a congeneric series of ligands. Specifically, we selected three targets with the largest datasets, *MCL1*, *HIF2A*, and *SYK*, containing 25, 37, and 43 unique compounds, respectively, to compare BALM+PEFT’s performance against AFE methods run as part of the OpenFE benchmark.^60^ Additionally, we evaluated docking performance on these targets using AutoDock Vina, by rescoring the 3D poses presented in the benchmark. Fig. 5 shows BALM+PEFT’s performance in zero-shot (a, b, c) and few-shot (d, e, f) modes for each target. In a zero-shot setting, the performance of the model is worse than both Vina and AFE, suggesting that although BALM+PEFT can predict binding free energy within a moderate error range, methods based on three-dimensional information can capture binding nuances more effectively, as recently pointed out by other authors.^5^ The few-shot setting, using ca. 20% of the data for each target (5 ligands for *MCL1*, 7 for *HIF2A*, and 8 for *SYK*), generally reduces the RMSE across all targets, although the rank correlation trends are very target-dependent, being slightly better for *SYK* and *MCL1*, but worse for *HIF2A*. Regardless of the zero or few-shot setting, the predictions obtained with BALM+PEFT cluster within a narrower binding affinity range than Vina or AFE, suggesting either limitations in distinguishing small R-group modifications to a common chemical scaffold or an unintended optimization of its loss function, optimizing for RMSE at the cost of capturing the full dynamic range of binding energies. Globally, these results hint that BALM+PEFT faces challenges in effectively ranking congeneric series, which complicates its potential as a direct alternative to AFE for lead optimization stages. This may stem from the limited few-shot data (5-8 ligands), preventing the model from capturing subtle R-group differences and potentially focusing more on improving RMSE during training without maintaining rank order. As we used a small subset of data in this study, future exploration should involve using larger datasets of congeneric series to better understand the impact of fine-tuning with data subsets of varying sizes. Additionally, implementing time-based splits of data collected over different periods could provide valuable insights by testing the model’s performance retrospectively. These steps may help in refining the model’s predictive capabilities and addressing its current limitations. However, the speed of inference and its higher accuracy with respect to methods such as molecular docking make the model an appealing alternative to popular re-ranking methods such as MM-GBSA,^61^ aimed at filtering out low-quality compounds and obtaining a fast answer in those situations in which speed may trump accuracy.

**Figure 5:**
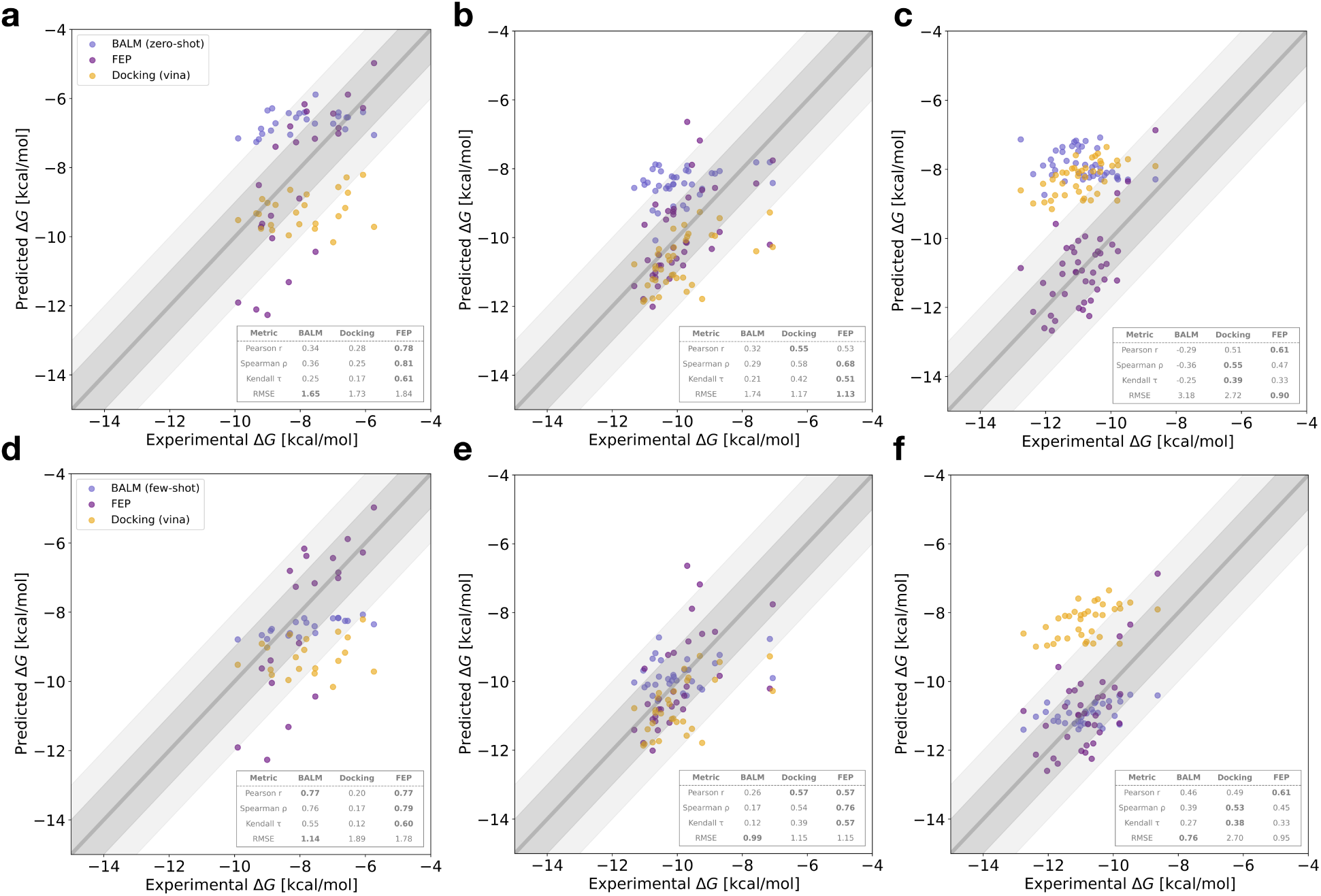
Comparing BALM, Alchemical Free Energy (AFE) predictions, and AutoDock Vina across three benchmark protein-ligand targets. The benchmark includes *MCL1* (25 ligands), *HIF2A* (37 ligands), and *SYK* (43 ligands). Panels **(a, b, c)** present the zero-shot predictions from BALM (blue) alongside AFE (yellow) and docking (orange) for each target. **(d, e, f)** illustrate the few-shot performance of BALM, fine-tuned with approximately 20% of the data. Each plot displays metrics for Pearson correlation, Spearman *ρ*, Kendall *τ*, and RMSE.

## Discussion

Drug discovery has many computational tools at its disposal in its early stages, from hit finding to lead-optimisation of a compound for a specific biomolecular target. Traditionally, these range from docking to simulation-based methods, and more recently include machine learning models such as AlphaFold3^62^ to aid in structure generation of the desired protein targets. One key objective for computer-aided drug discovery is to successfully predict affinities and other ADMET properties (Absorption, Distribution, Metabolism, Excretion, and Toxicity) of large molecular libraries, and later of a lead molecule, towards a given biomolecular target. For affinity predictions, there has historically been a trade-off between prediction speed (e.g., through a docking score processing millions of compounds at a time) and accuracy (e.g., through alchemical free energy methods, processing 100s of compounds at a time). More recently, machine learning methods for affinity predictions have been taking an increasingly central stage by complementing traditional methods. A current issue is that many machine-learning-based affinity prediction methods are not tested against single targets, or evaluated on target-specific metrics. Issues in the methodologies adopted to assess recent AI-driven docking approaches were highlighted by the case of DiffDock, where a careful re-assessment of its performance according to best practices yielded lower success rates than initially reported.^63,64^ In this work, we propose fair and robust evaluation strategies to test models’ performance and apply them to BALM, our new binding affinity prediction framework leveraging information on protein sequence and ligand SMILES.

As a sequence and SMILES-based model, BALM does not rely on structural data and can thus benefit from many large-language modelling tool developments such as parameter-efficient fine-tuning. In a target-specific setting, BALM with appropriate PEFT adapters outperforms baseline models on specific protein targets and can be further fine-tuned using a few-shot data scenario. This makes it a viable tool to be used alongside other computational methods for large-scale screening campaigns against individual targets. BALM is also fast and cost-effective, with few-shot learning on 100-300 compounds achieved in 10-20 minutes on typical GPU architectures such as Nvidia A100s. For targets without good crystallographic data available, BALM is a good alternative to traditional docking-based virtual screening methods. Current limitations preclude BALM from achieving alchemical free energy accuracy on congeneric ligand series often found in lead-optimisation campaigns. This can be addressed in multiple ways in the future looking at diverse ligand embeddings and other fine-tuning strategies. While its current performance does not provide a full replacement of traditional methods, BALM can be used alongside a docking campaign for consensus scoring and even replacement of computationally costly docking protocols where poor structural data is available.

Overall, BALM is likely to perform well in scenarios of single globular proteins with sufficient data availability for both fine-tuning and few-shot learning. In situations where proteins are homodimers, multimers, form part of a membrane, or have otherwise more complicated behavior (e.g., binding mediated by co-factors), BALM can complement more advanced strategies such as AFE. All of these situations constitute realistic drug discovery scenarios but may be captured well with language model-based approaches. For this reason, it is crucial that unbiased evaluation strategies are adopted to assess the performance of ML models. In this context, we show that looking at target-specific evaluation metrics and leak-free training of the framework is key to determining how well an ML model performs on a realistic affinity prediction task. In this work, we set a standard baseline on how ML-based affinity models should be evaluated. Only through the broad adoption of unbiased evaluation strategies such as those presented in our work, the community will obtain a real sense of the advance of machine-learning-based affinity predictions.

## Methods

### Model overview and objective

BALM is designed to predict protein-ligand binding affinity by taking ligand SMILES strings and protein sequences as the input. Our method begins by encoding the protein sequences and ligand SMILES by integrating protein and ligand language models. These language models trained on extensive ligand^40^ and protein databases^39^ encapsulate the chemical properties inherent in SMILES strings and physicochemical, functional, and evolutionary patterns in protein sequences. The encodings obtained from the language models are then projected through a linear layer to a shared latent space. The core idea behind BALM is to maximize the cosine similarity between protein-ligand embeddings for a binding interaction and minimize the same for non-binding interactions, thus learning protein-ligand interaction features. The model learns to understand the relationship between protein and ligand embeddings and quantifies it through a cosine similarity score, a proxy for the binding affinity score (pK*_d_*). The model learns by minimizing the mean-squared error (MSE) loss.

### Encoding and projecting protein and ligand features

We use ESM-2^32^ model for encoding protein features and ChemBERTa-2^33^ for encoding ligands. It is important to note that our focus is not to evaluate the performance of protein or ligand language models but to leverage their learned representations. The ESM-2 model is trained on sequences from the UniRef database^39^ with 65 million unique sequences using bidirectional transformer architecture.^65^ ESM-2 learns in an unsupervised manner through masked language modelling (MLM) objective, where 15% of amino acids are masked in the input sequence, and it is tasked with predicting these missing positions. We use the 150 million parameter model as it strikes a balance between computational efficiency and performance.

ChemBERTa-2 is based on the RoBERTa^66^ transformer model, using both MLM similar to ESM-2 by masking 15% tokens and multi-task regression (MTR) pretraining on 200 downstream molecular properties. We use the model trained on the largest dataset size using the MTR objective as it outperformed the other model configurations in the comparative study.^33^ For a given input protein sequence, *I_P_*, and ligand SMILES, *I_L_*, we encode and project to get embeddings of the same dimensions as shown below:

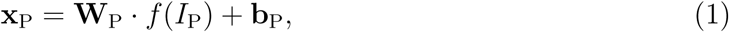

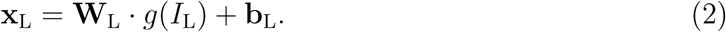

Here, 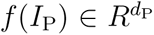 extracts the protein features using ESM-2 and similarly, 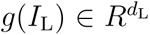 extracts the ligand features using ChemBERTa-2. We then transform them separately into **x**_P_, **x**_L_ ∈ *R^K^* using a single fully connected layer (with a ReLU activation). These layers are parameterized with weight matrices 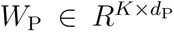 for proteins and 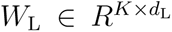 for ligands, and bias vectors **b**_P_, **b**_L_ ∈ *R^K^*. As the encodings derived from the ESM-2 and the ChemBERTa-2 model have different dimensions, 640 (150M model) and 384, respectively, we employ linear layers to map both sets of embeddings to a shared latent space of dimension *K* (256) suitable for similarity computations. Both ESM-2 and ChemBERTa-2 models are accessible via the Hugging face transformers library, ^67^ under the model names `facebook/esm2 t30 150M UR50D`, and `DeepChem/ChemBERTa-77M-MTR`, respectively.

### Training with cosine similarity and MSE

In the training phase of BALM, the cosine similarity metric is used to measure the affinity between the projected protein and ligand embeddings. BALM utilizes cosine similarity as the core metric for learning protein-ligand binding as a regression problem. BALM defines the binding score by computing the cosine similarity of the protein and ligand embeddings:

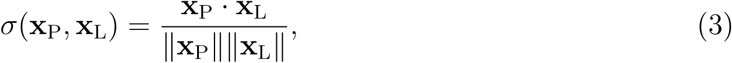

where **x**_P_ and **x**_L_ denote the embeddings of the protein and ligand, respectively. The cosine similarity *σ* ranges from −1 to 1, with values closer to 1 indicating a stronger binding (high pK*_d_*). 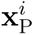 and 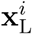 are the projected embeddings of the protein and ligand, respectively. BALM is trained by minimizing the MSE between predicted binding scores given by *σ*(**x**_P_, **x**_L_) and experimental affinities (*y_i_*) adjusted to the −1 to 1 scale :

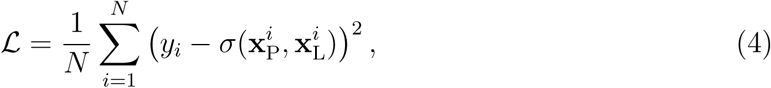

where *N* is the number of samples.

### Parameter-efficient fine-tuning on BALM

PEFT allows adapting pre-trained language models to new domains by fine-tuning a small subset of additional parameters while keeping the initial model parameters fixed. This reduces computational requirements and prevents catastrophic forgetting. Given a pre-trained model *T* with parameters *θ*, and a downstream task 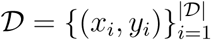, where (*x_i_, y_i_*) represents an input and ground-truth pair for task 𝒟, PEFT introduces a small set of trainable parameters Δ*θ*, where |Δ*θ*| ≪ |*θ*|. The objective is to adapt *θ* to task 𝒟 by optimizing:

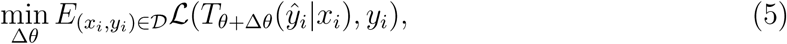

where *ŷ_i_* denotes the predicted affinities and ℒ denotes the MSE loss function reflecting the model’s performance on task 𝒟. Various PEFT methods have been proposed, for more details refer to the survey papers.^41,42^

In this work, we use the reparameterised and additive PEFT methods^41,42^ to fine-tune BALM’s protein (ESM-2) and ligand (ChemBERTa-2) language models for binding affinity prediction. The reparameterized PEFT methods we used in the study include Low-Rank Adaptation (LoRA),^68^ LoRA with Hadamard product (LoHa),^69^ and LoRA with Kronecker product (LoKr),^70^ applied to the key, query, and value matrices. An additive PEFT method, Infused Adapter by Inhibiting and Amplifying Inner Activations (IA^3^),^71^ is also studied for fine-tuning BALM. These methods are implemented using the Hugging face PeftModel class.^67,72^ We use four PEFT methods in our study, and these are discussed in detail in the Appendix **??**.

## Data for the study

To benchmark the performance of our models, we utilized several publicly available datasets, including BindingDB,^47,48^ LP-PDBBind,^55^ and other datasets specific to protein-ligand systems such as *USP7*, ^50^ *Mpro*,^51,52^ and three targets from protein-ligand free energy benchmark^8^ - *MCL1*, *HIF2A*, and *SYK*. These datasets encompass a wide range of binding affinity measurements and chemical diversity, enabling us to compare with docking and alchemical free energy methods. These datasets provide a comprehensive benchmark for strategically evaluating the machine learning model’s capability to predict binding affinity under various conditions, including zero-shot and few-shot scenarios. We release all the cleaned datasets used in our work as `BALM-Benchmark` on Hugging face.^67^

### BindingDB

dataset provides experimentally measured binding affinities of proteinligand interactions, we focus on the K*_d_* version^48^ due to inconsistencies in IC50 and K_i_ measures.^30^ This version contains 52,284 interactions with 10,665 ligand SMILES and 1,413 protein sequences. We filter out sequences with more than 1024 residues for computational efficiency with the ESM-2 model.^32^ The K_d_ values were transformed into pK_d_ for stable training, following previous works. ^24,29,35^ We were left with approximately 1100 targets and 48,000 interaction data after the filtering. Additionally, to avoid bias from assay limits (Fig. S1), we removed the top five most frequent limits. This reduced the dataset to about 25,000 interactions involving approximately 1,070 targets and 9,200 ligands. We used around 70% of these interaction data for training, 10% interactions for validation, and 20% interactions for testing. Four data splits (Random, Cold Target, Cold Drug, and Scaffold) were used to evaluate model generalizability. In the *random split*, commonly used in prior works,^24,29,35^ protein-ligand interaction pairs are randomly distributed across training, validation, and test sets, ensuring each subset is a statistically representative sample. The *cold target split* segregates protein targets into distinct groups for training, validation, and testing, allowing evaluation on unseen protein targets. Similarly, the *cold drug split* randomly allocates drugs to training, validation, and test sets, assigning all protein-ligand pairs linked to each ligand to the corresponding set, enabling performance evaluation on unseen drugs. Lastly, the *scaffold split* employs the Murcko scaffolds ^56^ concept, using the `MurckoScaffold` module from RDkit^57^ to bin drugs by their core scaffolds, ensuring no structural overlap across sets. Approximately 4570 scaffolds were used from the BindingDB dataset for this purpose.

### LP-PDBBind

dataset is derived from the publicly available PDBBind v2020^59^ dataset, a curated set of approximately 20,000 protein-ligand complex structures (3D) with experimentally measured binding affinities. The dataset has been reorganized to minimize sequence, structural interaction patterns, and chemical similarity across training, validation, and test splits.^55^ It has also been cleaned to remove covalent-bound ligand-protein complexes, ligands with rare atomic elements, and structures with steric clashes while maintaining consistency in reported binding free energies. Recently, Jores et al.^73^ provided more detailed analysis of the LP-PDBBind’s data splits. Furthermore, the dataset has been categorized based on the quality of structures into Clean Levels 1, 2, and 3, consisting of 14,324, 7,985, and 4,404 entries, respectively. We utilize CL1 for training, while CL2 is employed for validation and testing due to its higher reliability, as suggested by Li et al.^55^

### USP7

dataset is curated by Shen et al.,^50^ evaluates inhibitors targeting the ubiquitin-specific protease 7 (*USP7*). The dataset consists of over 4000 ligands with associated binding affinities collected from ChEMBL.^74^ After processing, the final dataset comprises 1,799 unique ligands with measured experimental affinities, represented as IC50 values. These values were then transformed into pIC50 values for uniformity and stability in training.

### Mpro

dataset is derived from the COVID Moonshot project focusing on inhibitors of the SARS-CoV-2 main protease (*Mpro*),^51,52^ includes data from multiple design sprints. We filtered to remove assay limits, and the cleaned dataset contains 2,062 unique ligands with experimentally determined IC50 values. We converted IC50 values to pIC50 values similar to the *USP7* target. Further details about the data curation can be found here.^51,52^

### Protein-ligand free energy benchmark

curated by Hahn et al.^8^ contains about 21 targets for benchmarking alchemical free energy calculations. We selected three targets, *MCL1*, *HIF2A*, and *SYK*, to benchmark the machine learning model performance as compared to the alchemical energy calculations,^60^ provided through the OpenFE (open free energy) consortium. The *HIF2A* dataset contains 37 ligands, the *MCL1* dataset includes 25 ligands, and the *SYK* dataset consists of 43 unique ligands.

## Acknowledgement

RG and APG were supported by the United Kingdom Research and Innovation (grant EP/S02431X/1), UKRI Centre for Doctoral Training in Biomedical AI at the University of Edinburgh, School of Informatics. RG was also supported by Exscientia Plc, Oxford, now Recusrion. Experiments from this work are conducted mainly on the Edinburgh International Data Facility^1^ and supported by the Data-Driven Innovation Programme at the University of Edinburgh.

## Supporting Information Available

Supporting Information Available: Dataset analysis, additional experimental results and methods used are provided.

## Data and Code Availability

All datasets curated for this study are publicly available via the Hugging Face BALM-Benchmark collection. The datasets can be accessed at https://huggingface.co/datasets/BALM/BALM-benchmark, and the pre-trained models are accessible on Hugging Face at https://huggingface.co/BALM. All the code containing scripts for data processing, model training, and evaluation is publicly accessible on GitHub at https://github.com/meyresearch/BALM

## Footnotes

1 https://edinburgh-international-data-facility.ed.ac.uk/

## References

(1) Lyu, J.; Wang, S.; Balius, T. E.; Singh, I.; Levit, A.; Moroz, Y. Y.; O’Meara, M. J.; Che, T.; Algaa, E.; Tolmachova, K.; Tolmachev, A. A. Ultra-large library docking for discovering new chemotypes. Nature 2019, 566, 224–229.

(2) Stanzione, F.; Giangreco, I.; Cole, J. C. Use of molecular docking computational tools in drug discovery. Prog. Med. Chem. 2021, 60, 273–343.

(3) Shoichet, B. K. Virtual screening of chemical libraries. Nature 2004, 432, 862–865.

(4) Beroza, P.; Crawford, J. J.; Ganichkin, O.; Gendelev, L.; Harris, S. F.; Klein, R.; Miu, A.; Steinbacher, S.; Klingler, F.-M.; Lemmen, C. Chemical space docking enables large-scale structure-based virtual screening to discover ROCK1 kinase inhibitors. Nat. Commun. 2022, 13, 6447.

(5) Errington, D.; Schneider, C.; Bouysset, C.; Dreyer, F. A. Assessing interaction recovery of predicted protein-ligand poses. arXiv:2409.20227 2024,

(6) Robo, M. T.; Hayes, R. L.; Ding, X.; Pulawski, B.; Vilseck, J. Z. Fast free energy estimates from *λ*-dynamics with bias-updated Gibbs sampling. Nat. Commun. 2023, 14, 8515.

(7) Mey, A. S.; Allen, B. K.; Macdonald, H. E. B.; Chodera, J. D.; Hahn, D. F.; Kuhn, M.; Michel, J.; Mobley, D. L.; Naden, L. N.; Prasad, S.; Rizzi, A.; Scheen, J.; Shirts, M. R.; Tresadern, G.; Xu, H. Best Practices for Alchemical Free Energy Calculations. Living J. Mol. Sci. 2020, 2, 18378.

(8) Hahn, D. F.; Bayly, C. I.; Boby, M. L.; Macdonald, H. E. B.; Chodera, J. D.; Gapsys, V.; Mey, A. S.; Mobley, D. L.; Benito, L. P.; Schindler, C. E.; Tresadern, G.; Warren, G. L. Best practices for constructing, preparing, and evaluating protein-ligand binding affinity benchmarks. Living J. Mol. Sci. 2022, 4, 1497–1497.

(9) Sadybekov, A. V.; Katritch, V. Computational approaches streamlining drug discovery. Nature 2023, 616, 673–685.

(10) Grygorenko, O. O.; Radchenko, D. S.; Dziuba, I.; Chuprina, A.; Gubina, K. E.; Moroz, Y. S. Generating multibillion chemical space of readily accessible screening compounds. Iscience 2020, 23.

(11) Hadfield, T. E.; Scantlebury, J.; Deane, C. M. Exploring the ability of machine learning-based virtual screening models to identify the functional groups responsible for binding. J. Cheminformatics 2023, 15, 84.

(12) Zhang, Y.; Li, S.; Meng, K.; Sun, S. Machine Learning for Sequence and Structure-Based Protein–Ligand Interaction Prediction. J. Chem. Inf. Model. 2024,

(13) Kimber, T. B.; Chen, Y.; Volkamer, A. Deep learning in virtual screening: recent applications and developments. Int. J. Mol. Sci. 2021, 22, 4435.

(14) Smer-Barreto, V.; Quintanilla, A.; Elliott, R. J. R.; Dawson, J. C.; Sun, J.; Campa, V. M.; Lorente-Macías, Á.; Unciti-Broceta, A.; Carragher, N. O.; Acosta, J. C.; others Discovery of senolytics using machine learning. Nat. Commun. 2023, 14, 3445.

(15) Rodriguez, S.; Hug, C.; Todorov, P.; Moret, N.; Boswell, S. A.; Evans, K.; Zhou, G.; Johnson, N. T.; Hyman, B. T.; Sorger, P. K.; others Machine learning identifies candidates for drug repurposing in Alzheimer’s disease. Nat. Commun. 2021, 12, 1033.

(16) Gusev, F.; Gutkin, E.; Kurnikova, M. G.; Isayev, O. Active learning guided drug design Lead optimization based on relative binding free energy modeling. J. Chem. Inf. Model. 2023, 63, 583–594.

(17) Thompson, J.; Walters, W. P.; Feng, J. A.; Pabon, N. A.; Xu, H.; Goldman, B. B.; Moustakas, D.; Schmidt, M.; York, F. Optimizing active learning for free energy calculations. Artif. Intell. Life Sci. 2022, 2, 100050.

(18) Gorantla, R.; Kubincova, A.; Suutari, B.; Cossins, B. P.; Mey, A. S. Benchmarking active learning protocols for ligand binding affinity prediction. J. Chem. Inf. Model. 2024, 64, 1955–1965.

(19) Huang, L.; Xu, T.; Yu, Y.; Zhao, P.; Chen, X.; Han, J.; Xie, Z.; Li, H.; Zhong, W.; Wong, K.-C.; others A dual diffusion model enables 3D molecule generation and lead optimization based on target pockets. Nat. Commun. 2024, 15, 2657.

(20) Anstine, D. M.; Isayev, O. Generative models as an emerging paradigm in the chemical sciences. J. Am. Chem. Soc. 2023, 145, 8736–8750.

(21) Sattari, K.; Li, D.; Kalita, B.; Xie, Y.; Lighvan, F. B.; Isayev, O.; Lin, J. De novo molecule design towards biased properties via a deep generative framework and iterative transfer learning. Digit. Discov. 2024,

(22) Runcie, N. T.; Mey, A. S. SILVR: Guided diffusion for molecule generation. J. Chem. Inf. Model. 2023, 63, 5996–6005.

(23) Chatterjee, A.; Walters, R.; Shafi, Z.; Ahmed, O. S.; Sebek, M.; Gysi, D.; Yu, R.; Eliassi-Rad, T.; Barabási, A.-L.; Menichetti, G. Improving the generalizability of protein-ligand binding predictions with AI-Bind. Nat. Commun. 2023, 14, 1989.

(24) Öztürk, H.; Özgür, A.; Ozkirimli, E. DeepDTA: deep drug–target binding affinity prediction. Bioinformatics 2018, 34, 821–829.

(25) Jiménez, J.; Skalic, M.; Martinez-Rosell, G.; De Fabritiis, G. K deep: protein–ligand absolute binding affinity prediction via 3d-convolutional neural networks. J. Chem. Inf. Model. 2018, 58, 287–296.

(26) Shen, L.; Feng, H.; Li, F.; Lei, F.; Wu, J.; Wei, G.-W. Knot data analysis using multiscale Gauss link integral. Proc. Natl. Acad. Sci. USA 2024, 121, e2408431121.

(27) Wang, R.; Fang, X.; Lu, Y.; Wang, S. The PDBbind database: Collection of binding affinities for proteinligand complexes with known three-dimensional structures. J. Med. Chem. 2004, 47, 2977–2980.

(28) Volkov, M.; Turk, J.-A.; Drizard, N.; Martin, N.; Hoffmann, B.; Gaston-Mathé, Y.; Rognan, D. On the frustration to predict binding affinities from protein–ligand structures with deep neural networks. J. Med. Chem. 2022, 65, 7946–7958.

(29) Gorantla, R.; Kubincova, A.; Weiße, A. Y.; Mey, A. S. From Proteins to Ligands: Decoding Deep Learning Methods for Binding Affinity Prediction. J. Chem. Inf. Model. 2024, 64, 2496–2507.

(30) Landrum, G. A.; Riniker, S. Combining IC50 or K i Values from Different Sources Is a Source of Significant Noise. J. Chem. Inf. Model. 2024,

(31) Backenköhler, M.; Groß, J.; Wolf, V.; Volkamer, A. Guided docking as a data generation approach facilitates structure-based machine learning on kinases. J. Chem. Inf. Model. 2024, 64, 4009–4020.

(32) Lin, Z.; Akin, H.; Rao, R.; Hie, B.; Zhu, Z.; Lu, W.; Smetanin, N.; Verkuil, R.; Kabeli, O.; Shmueli, Y.; others Evolutionary-scale prediction of atomic-level protein structure with a language model. Science 2023, 379, 1123–1130.

(33) Ahmad, W.; Simon, E.; Chithrananda, S.; Grand, G.; Ramsundar, B. Chemberta-2: Towards chemical foundation models. arXiv:2209.01712 2022,

(34) Nguyen, T.; Le, H.; Quinn, T. P.; Nguyen, T.; Le, T. D.; Venkatesh, S. GraphDTA: Predicting drug–target binding affinity with graph neural networks. Bioinformatics 2021, 37, 1140–1147.

(35) Jiang, M.; Li, Z.; Zhang, S.; Wang, S.; Wang, X.; Yuan, Q.; Wei, Z. Drug–target affinity prediction using graph neural network and contact maps. RSC Adv. 2020, 10, 20701–20712.

(36) Luo, D.; Liu, D.; Qu, X.; Dong, L.; Wang, B. Enhancing Generalizability in Protein– Ligand Binding Affinity Prediction with Multimodal Contrastive Learning. J. Chem. Inf. Model. 2024,

(37) Kaufman, B.; Williams, E. C.; Underkoffler, C.; Pederson, R.; Mardirossian, N.; Watson, I.; Parkhill, J. Coati: Multimodal contrastive pretraining for representing and traversing chemical space. J. Chem. Inf. Model. 2024, 64, 1145–1157.

(38) Singh, R.; Sledzieski, S.; Bryson, B.; Cowen, L.; Berger, B. Contrastive learning in protein language space predicts interactions between drugs and protein targets. Proc. Natl. Acad. Sci. U.S.A. 2023, 120, e2220778120.

(39) Suzek, B. E.; Wang, Y.; Huang, H.; McGarvey, P. B.; Wu, C. H.; Consortium, U. UniRef clusters: a comprehensive and scalable alternative for improving sequence similarity searches. Bioinformatics 2015, 31, 926–932.

(40) Kim, S.; Chen, J.; Cheng, T.; Gindulyte, A.; He, J.; He, S.; Li, Q.; Shoemaker, B. A.; Thiessen, P. A.; Yu, B.; Zaslavsky, L.; Zhang, J.; E Bolton, E. PubChem 2019 update: improved access to chemical data. Nucleic Acids Res. 2019, 47, D1102–D1109.

(41) Han, Z.; Gao, C.; Liu, J.; Zhang, S. Q. Parameter-Efficient Fine-Tuning for Large Models: A Comprehensive Survey. arXiv:2403.14608 2024,

(42) Xu, L.; Xie, H.; Qin, S.-Z. J.; Tao, X.; Wang, F. L. Parameter-efficient fine-tuning methods for pretrained language models: A critical review and assessment. arXiv:2312.12148 2023,

(43) Sultan, A.; Sieg, J.; Mathea, M.; Volkamer, A. Transformers for molecular property prediction: Lessons learned from the past five years. J. Chem. Inf. Model. 2024, 64, 6259–6280.

(44) Dutt, R.; Ericsson, L.; Sanchez, P.; Tsaftaris, S. A.; Hospedales, T. Parameter-efficient fine-tuning for medical image analysis: The missed opportunity. arXiv:2305.08252 2023,

(45) Gema, A.; Daines, L.; Minervini, P.; Alex, B. Parameter-efficient fine-tuning of llama for the clinical domain. arXiv:2307.03042 2023,

(46) Gema, A. P.; Hong, G.; Minervini, P.; Daines, L.; Alex, B. Edinburgh Clinical NLP at SemEval-2024 Task 2: Fine-tune your model unless you have access to GPT-4. arXiv:2404.00484 2024,

(47) Gilson, M. K.; Liu, T.; Baitaluk, M.; Nicola, G.; Hwang, L.; Chong, J. BindingDB in 2015: a public database for medicinal chemistry, computational chemistry and systems pharmacology. Nucleic Acids Res. 2016, 44, D1045–D1053.

(48) Huang, K.; Fu, T.; Gao, W.; Zhao, Y.; Roohani, Y.; Leskovec, J.; Coley, C. W.; Xiao, C.; Sun, J.; Zitnik, M. Therapeutics data commons: Machine learning datasets and tasks for drug discovery and development. arXiv:2102.09548 2021,

(49) Fisher, R. Frequency Distribution of the Values of the Correlation Coefficient in Samples from an Indefinitely Large Population. Biometrika 1915, 10, 507–521.

(50) Shen, W.-f.; Tang, H.-w.; Li, J.-b.; Li, X.; Chen, S. Multimodal data fusion for supervised learning-based identification of USP7 inhibitors: a systematic comparison. J. Cheminform. 2023, 15, 1–16.

(51) Achdout, H.; Aimon, A.; Bar-David, E.; Morris, G. COVID moonshot: open science discovery of SARS-CoV-2 main protease inhibitors by combining crowdsourcing, highthroughput experiments, computational simulations, and machine learning. BioRxiv 2020,

(52) Boby, M. L.; Fearon, D.; Ferla, M.; Filep, M.; Koekemoer, L.; Robinson, M. C.; Consortium, T. C. M.; Chodera, J. D.; Lee, A. A.; London, N.; von Delft, A.; von Delft, F. Open science discovery of potent noncovalent SARS-CoV-2 main protease inhibitors. Science 2023, 382, eabo7201.

(53) Trott, O.; Olson, A. J. AutoDock Vina: improving the speed and accuracy of docking with a new scoring function, efficient optimization, and multithreading. J. Comput. Chem. 2010, 31, 455–461.

(54) Ruiz-Carmona, S.; Alvarez-Garcia, D.; Foloppe, N.; Gago, F.; Gervais, G.; Irwin, J.; Sverrisson, F.; Tounge, B.; Tresadern, G.; Morley, S. rDock: A Fast, Versatile and Open Source Program for Docking Ligands to Proteins and Nucleic Acids. PLoS Comput. Biol. 2014, 10, e1003571.

(55) Li, J.; Guan, X.; Zhang, O.; Sun, K.; Wang, Y.; Bagni, D.; Head-Gordon, T. Leak Proof PDBBind: A Reorganized Dataset of Protein-Ligand Complexes for More Generalizable Binding Affinity Prediction. arXiv:2308.09639 2023,

(56) Bemis, G. W.; Murcko, M. A. The properties of known drugs. 1. Molecular frameworks. J. Med. Chem. 1996, 39, 2887–2893.

(57) Bento, A. P.; Hersey, A.; Félix, E.; Landrum, G.; Gaulton, A.; Atkinson, F.; Bellis, L. J.; De Veij, M.; Leach, A. R. An open source chemical structure curation pipeline using RDKit. J. Cheminform. 2020, 12, 1–16.

(58) Deringer, V. L.; Bartók, A. P.; Bernstein, N.; Wilkins, D. M.; Ceriotti, M.; Csányi, G. Gaussian process regression for materials and molecules. Chem. Rev. 2021, 121, 10073–10141.

(59) Liu, Z.; Li, Y.; Han, L.; Li, J.; Liu, J.; Zhao, Z.; Nie, W.; Liu, Y.; Wang, R. PDB-wide collection of binding data: current status of the PDBbind database. Bioinformatics 2015, 31, 405–412.

(60) Baumann, H. M.; Henry, M. M.; Ries, B.; Swenson, D. W. H.; Eastwood, J. R. B.; Gowers, R. J.; Alibay, I. Openfe 1.0rc Release: Benchmarking Results. 2024; 10.5281/zenodo.13959654, Accessed: October 21, 2024.

(61) Genheden, S.; Ryde, U. The MM/PBSA and MM/GBSA methods to estimate ligand-binding affinities. Expert Opin. Drug Discov. 2015, 10, 449–461.

(62) Abramson, J.; Adler, J.; Dunger, J.; Evans, R.; Green, T.; Pritzel, A.; Ronneberger, O.; Willmore, L.; Ballard, A. J.; Bambrick, J.; others Accurate structure prediction of biomolecular interactions with AlphaFold 3. Nature 2024, 630, 493–500.

(63 ) Corso, G.; Stärk, H.; Jing, B.; Barzilay, R.; Jaakkola, T. Diffdock: Diffusion steps, twists, and turns for molecular docking. arXiv:2210.01776 2022,

(64) Jain, A. N.; Cleves, A. E.; Walters, W. P. Deep-Learning Based Docking Methods: Fair Comparisons to Conventional Docking Workflows. arXiv:2412.02889 2024,

(65) Devlin, J.; Chang, M.-W.; Lee, K.; Toutanova, K. Bert: Pre-training of deep bidirectional transformers for language understanding. arXiv:1810.04805 2018,

(66) Liu, Y.; Ott, M.; Goyal, N.; Du, J.; Joshi, M.; Chen, D.; Levy, O.; Lewis, M.; Zettlemoyer, L.; Stoyanov, V. Roberta: A robustly optimized bert pretraining approach. arXiv:1907.11692 2019,

(67) Wolf, T.; Debut, L.; Sanh, V.; Chaumond, J.; Delangue, C.; Moi, A.; Cistac, P.; Rault, T.; Louf, R.; Funtowicz, M.; others Huggingface’s transformers: State-of-the-art natural language processing. arXiv:1910.03771 2019,

(68) Hu, E. J.; Shen, Y.; Wallis, P.; Allen-Zhu, Z.; Li, Y.; Wang, S.; Wang, L.; Chen, W. Lora: Low-rank adaptation of large language models. arXiv:2106.09685 2021,

(69) Yeh, S.-Y.; Hsieh, Y.-G.; Gao, Z.; Yang, B. B.; Oh, G.; Gong, Y. Navigating text-to-image customization: From lycoris fine-tuning to model evaluation. arXiv:2309.14859 2023,

(70) He, X.; Li, C.; Zhang, P.; Yang, J.; Wang, X. E. Parameter-Efficient Model Adaptation for Vision Transformers. Thirty-Seventh AAAI Conference on Artificial Intelligence, AAAI 2023, Thirty-Fifth Conference on Innovative Applications of Artificial Intelligence, IAAI 2023, Thirteenth Symposium on Educational Advances in Artificial Intelligence, EAAI 2023, Washington, DC, USA, February 7-14, 2023. 2023; pp 817–825.

(71) Liu, H.; Tam, D.; Muqeeth, M.; Mohta, J.; Huang, T.; Bansal, M.; Raffel, C. A. Few-shot parameter-efficient fine-tuning is better and cheaper than in-context learning. Adv. Neural Inf. Process. Syst. 2022, 35, 1950–1965.

(72) Mangrulkar, S.; Gugger, S.; Debut, L.; Belkada, Y.; Paul, S.; Bossan, B. PEFT: State-of-the-art Parameter-Efficient Fine-Tuning methods. https://github.com/huggingface/peft, 2022.

(73) Joeres, R.; Blumenthal, D. B.; Kalinina, O. V. DataSAIL: Data Splitting Against Information Leakage. bioRxiv 2023, 2023–11.

(74) Mendez, D. et al. ChEMBL: towards direct deposition of bioassay data. Nucleic Acids Res. 2019, 47, D930–D940.

