## Supplementary Information for "Learning Binding Affinities via Fine-tuning of Protein and Ligand Language Models"

‖*Independent Researcher, 37027, United States*

### Additional Results

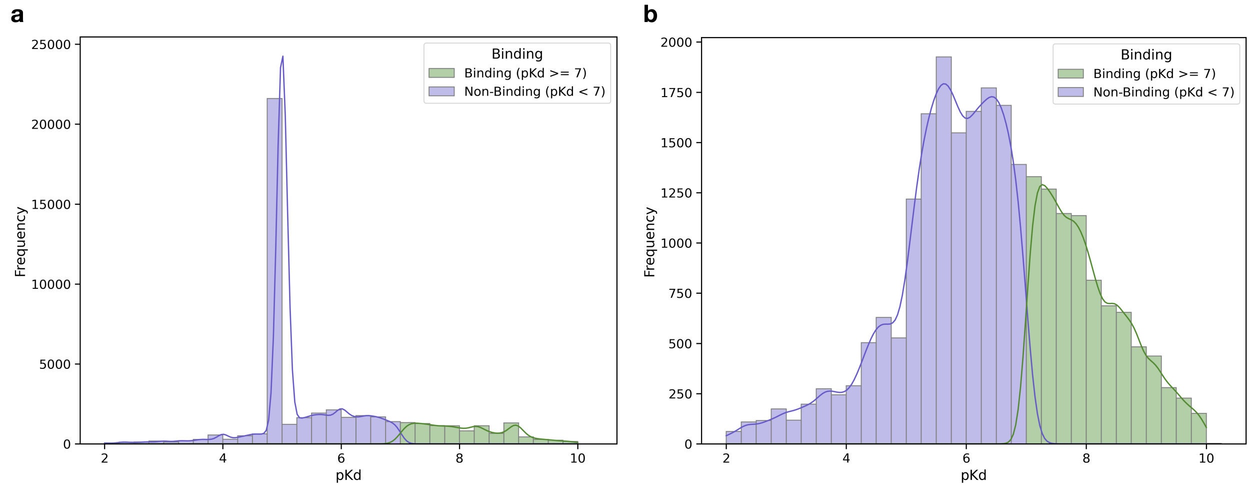


Figure S1: **BindingDB distribution of $pK_d$ values before and after removing assay limits.** **(a)** The distribution of binding affinities ($pK_d$) in the original BindingDB dataset shows significant skewness, primarily caused by the dominance of interactions with assay limits, where the most frequent value ($pK_d = 4.99$) accounts for a large portion of the data. **(b)** After removing the top five most frequent assay limits, the dataset demonstrates a more balanced distribution of binding affinities, providing a better foundation for model training and reducing prediction bias.

##### BindingDB data distribution

**Parameter-efficient fine-tuning significantly improves BALM's performance** For both ligand and protein-only fine-tuning, we tested various PEFT methods and ranks. LoHa and LoKr showed improvement in performance across all ranks as compared to the no fine-tuning model, while LoRA showed improvement with rank 8. However, the performance dropped at higher ranks. LoHa seemed to be stable, with rank 16 giving the best improvement of approximately 9.4%, showing a significant improvement over the BALM model without fine-tuning. Other ranks for LoHa also provided notable improvements, achieving around 8.7% (rank 8) and 8.3% (rank 32). IA3, while beneficial, offered a moderate gain of about 5.0%. In protein-only fine-tuning, LoKr seemed to be consistent across all ranks, with the highest improvement of approximately 18.2% at rank 8. Other ranks for LoKr also showed significant improvements, achieving gains of approximately 17.4% (ranks 16 and 32). IA3 and LoHa also demonstrated substantial improvements, with LoHa (rank 8) achieving

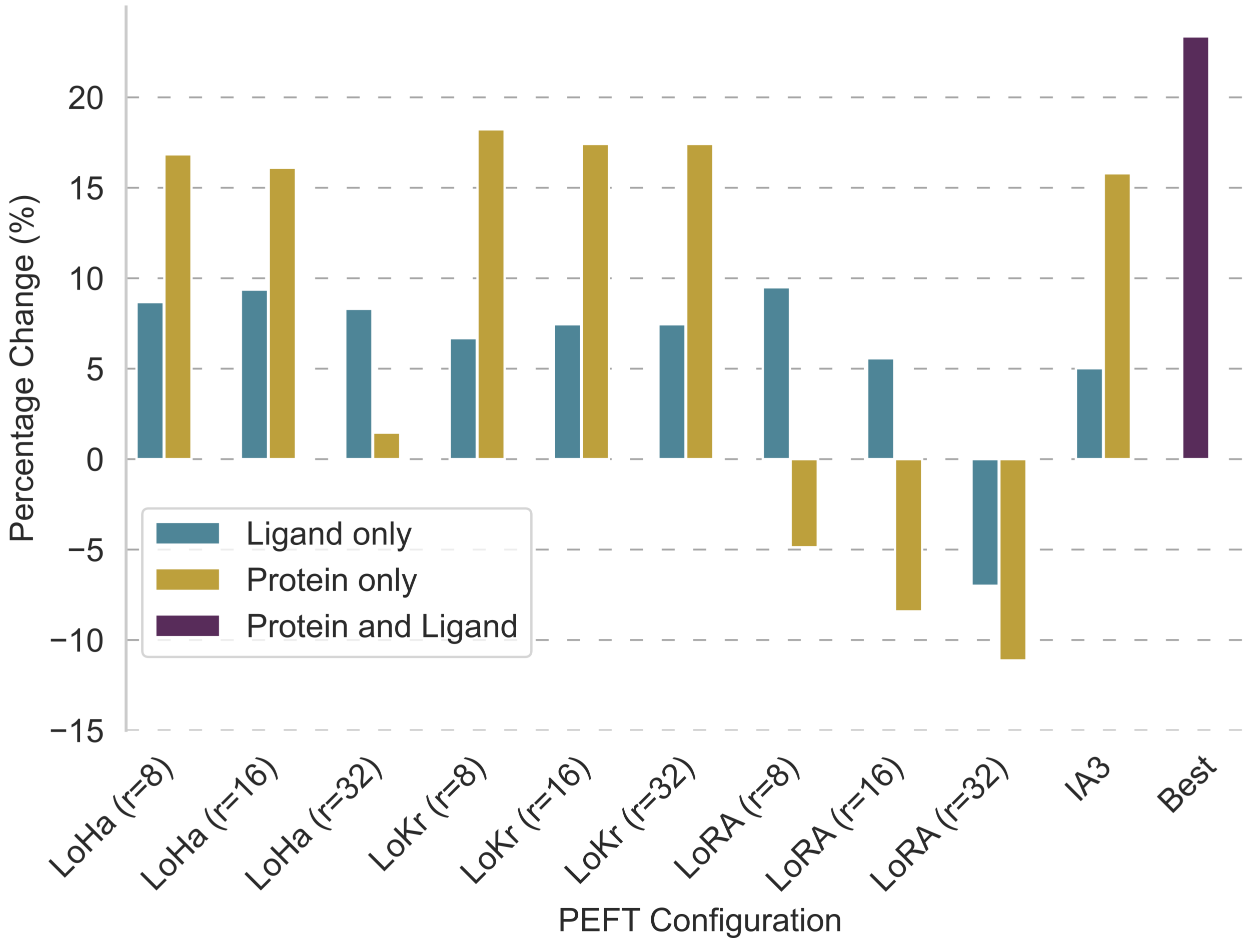


Figure S2: **Comparative analysis of parameter-efficient fine-tuning (PEFT) methods for protein and ligand language models.** This study evaluates the performance of various PEFT methods, specifically LoRA, LoHa, LoKr, and $IA^3$, in fine-tuning protein and ligand language models within the BALM framework. The performance of these methods is measured using Pearson correlation and compared against BALM without fine-tuning. Initially, we fine-tune protein and ligand language models separately in the BALM framework. Subsequently, the most effective fine-tuning methods (denoted as Best in the plot) for protein (LoKr) and ligand (LoHa) models are applied to assess their combined impact on the performance of BALM+PEFT.

around 16.8% gain and LoHa (rank 16) showing a gain of 16.1%. In contrast, LoRA showed a drop in performance. Finally, we combined the best-performing fine-tuning methods, LoHa for ligands and LoKr for proteins, and chose rank 16 for both to keep it consistent to create the BALM+PEFT variant. This combination yielded a performance gain of around 23.4% compared to BALM with no fine-tuning.

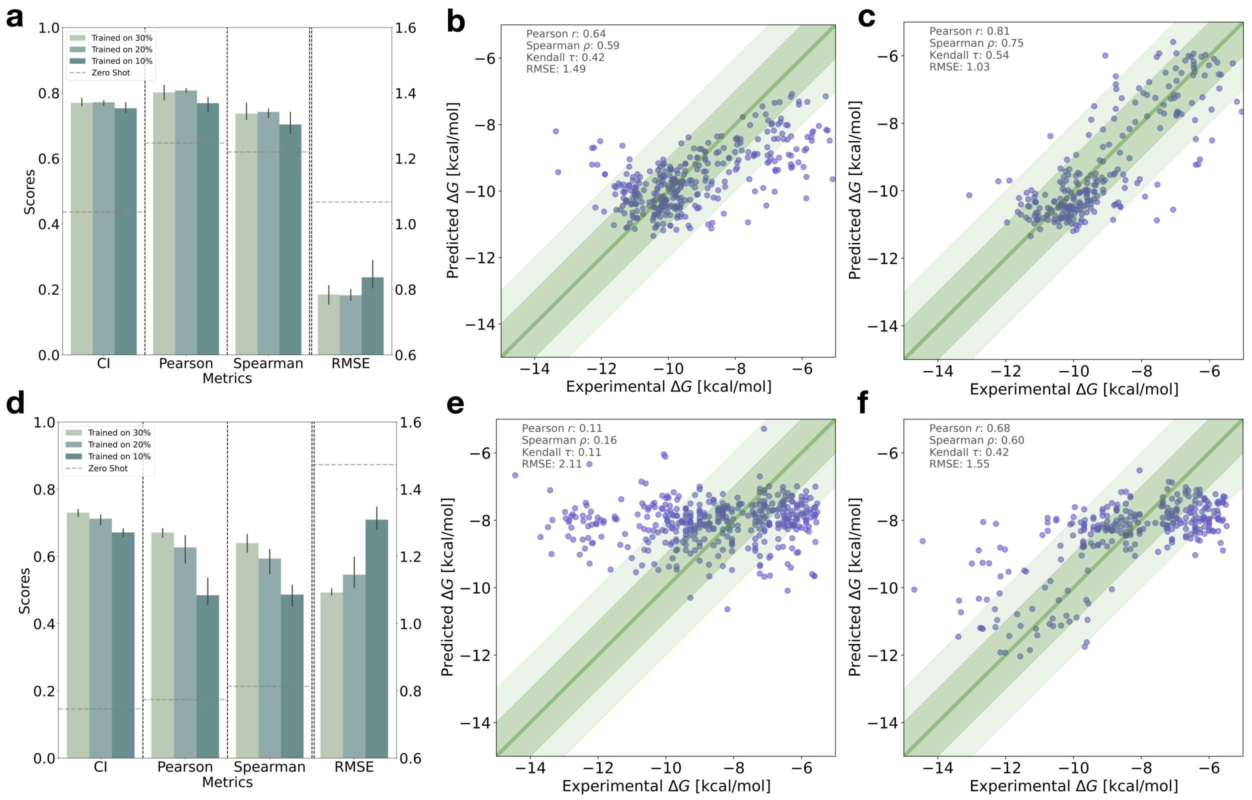


Figure S3: **Zero-shot and few-shot performance of the BALM+PEFT model on *Mpro* and *USP7* targets. (a, d)** Performance comparison of the pre-trained BALM+PEFT model (zero-shot) and few-shot fine-tuning using 10%, 20%, and 30% of experimental data. Testing is performed on the remaining *USP7* (a) and *Mpro* (d) data. The model is fine-tuned by retraining only the projection layer, and the data is split randomly with three different seeds. Error bars indicate the standard deviation across different splits, and performance metrics are reported for Concordance Index (CI), Pearson correlation, Spearman rank correlation, and Root Mean Squared Error (RMSE). **(b, e)** Scatter plots showing zero-shot model predictions for *USP7* (b) and *Mpro* (e) targets. Experimental $\Delta G$ values (kcal/mol) are on the x-axis and predicted $\Delta G$ values are on the y-axis. Only 20% of the test set (selected randomly) is shown for readability. **(c, f)** Scatter plots showing few-shot model performance on 20% of the training data for *USP7* (c) and *Mpro* (f) targets, with experimental $\Delta G$ on the x-axis and predicted $\Delta G$ on the y-axis. Metrics for Pearson $R$, Spearman $\rho$, Kendall $\tau$, and RMSE are displayed in the top-left corner of each scatter plot. Only 20% of the test set (selected randomly) is shown for readability.

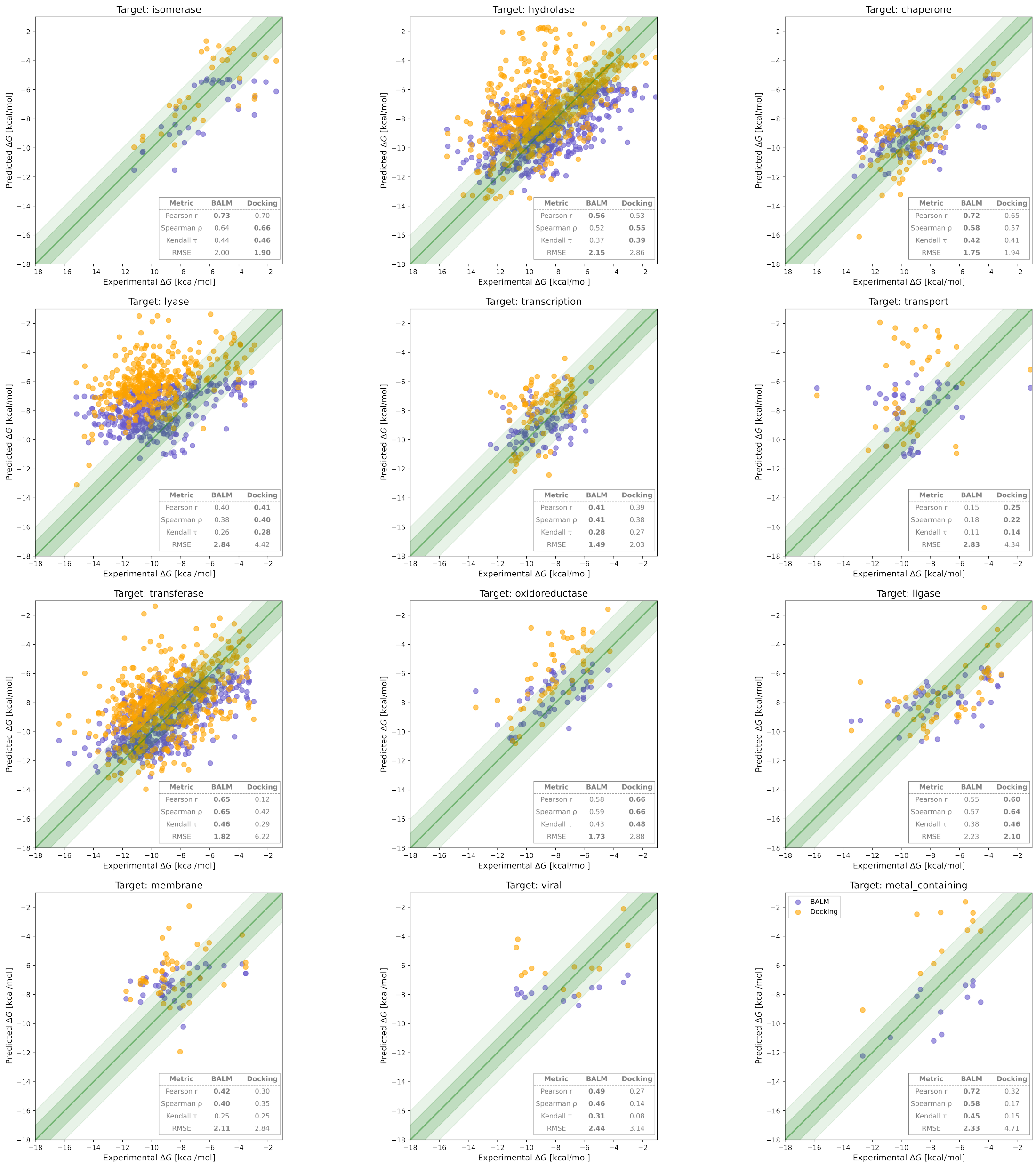


Figure S4: **Comparing BALM's (purple) and Autodock Vina's (orange) performance across different target families in the LP-PDBBind dataset.** Scatter plots show the predicted versus experimental binding affinities ($\Delta G$) for various target types in the zero-shot setting using the BALM+PEFT model. Target families include a wide range of protein classes, such as isomerases, hydrolases, and membrane proteins. Pearson correlation ($r$), Kendall $\tau$ Spearman rank correlation ($\rho$), and RMSE are shown for each target type.

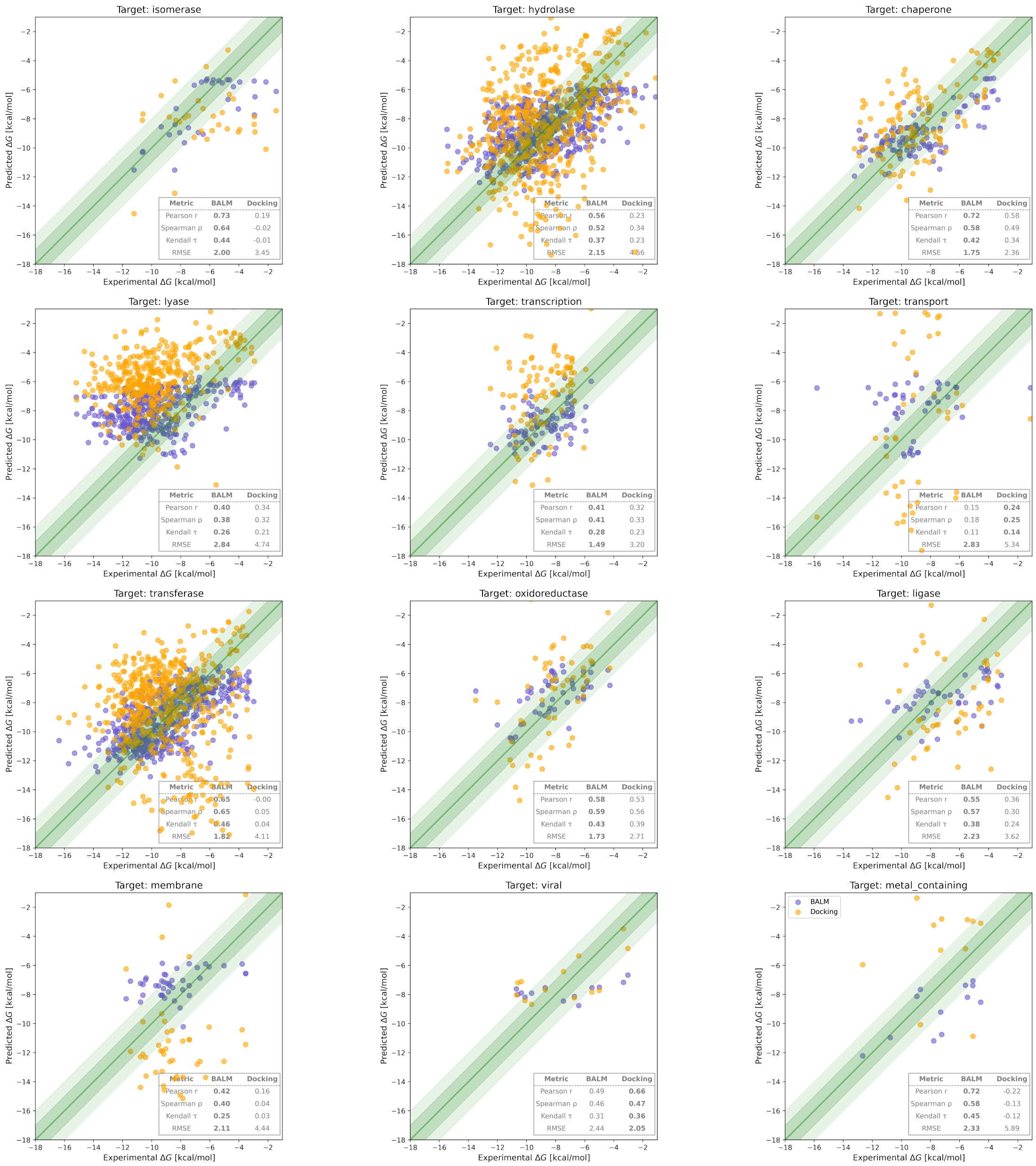

Figure S5: **Comparing BALM's (purple) and rDock (orange) performance across different target families in the LP-PDBBind dataset.** Scatter plots show the predicted versus experimental binding affinities ($\Delta G$) for various target types in the zero-shot setting using the BALM+PEFT model. Target families include a wide range of protein classes, such as isomerases, hydrolases, and membrane proteins. Pearson correlation ($r$), Kendall $\tau$ Spearman rank correlation ($\rho$), and RMSE are shown for each target type.

### Additional Methods

#### PEFT techniques

The strategies for fine-tuning the protein and ligand language models used in the study are discussed below.

- **LoRA** introduces $\Delta\mathbf{W} = \mathbf{BA}$, where $\mathbf{B} \in R^{p\times r}$ and $\mathbf{A} \in R^{r\times q}$ are trainable low-rank matrices, with $r \ll \min(p, q)$. This modification is added to the pre-trained weight $\mathbf{W}_0$, yielding an updated forward pass:

$$\mathbf{h}' = \mathbf{W}_0\mathbf{h} + \mathbf{b} + \gamma(\mathbf{BA})\mathbf{h}, \tag{1}$$

  where $\mathbf{h}$ represents the input and $\mathbf{h}'$ represents the output after the forward pass using the updated weights. $\gamma$ is a scaling factor to control the initialization of $\mathbf{B}$ and $\mathbf{A}$, balancing pre-trained knowledge and new adaptations, with $\gamma = \alpha/r$ for some defined $\alpha$, and $r$ is the rank of the matrix.

- **LoHa** uses $\Delta\mathbf{W} = (\mathbf{B}_1\mathbf{A}_1) \odot (\mathbf{B}_2\mathbf{A}_2)$, enhancing the rank of the update matrix and thus the fine-tuning capacity without significantly increasing parameter count. The forward pass becomes:

$$\mathbf{h}' = \mathbf{W}_0\mathbf{h} + \mathbf{b} + \gamma[(\mathbf{B}_1\mathbf{A}_1) \odot (\mathbf{B}_2\mathbf{A}_2)]\mathbf{h}, \tag{2}$$

  where $\odot$ denotes the Hadamard (element-wise) product.

- **LoKr** employs $\Delta\mathbf{W} = \mathbf{C} \otimes (\mathbf{BA})$, leveraging the multiplicative rank property of Kronecker products to extend beyond low-rank constraints while maintaining parameter efficiency. The forward pass is updated as:

$$\mathbf{h}' = \mathbf{W}_0\mathbf{h} + \mathbf{b} + \gamma(\mathbf{C} \otimes (\mathbf{BA}))\mathbf{h}, \tag{3}$$

where $\otimes$ denotes the Kronecker product.

- **IA**$^3$ method rescales the inner activations of the self-attention mechanism in transformer layers using learned vectors. IA$^3$ introduces three learnable rescaling vectors $\mathbf{l}_k \in R^{d_k}$, $\mathbf{l}_v \in R^{d_v}$, and $\mathbf{l}_f \in R^{d_{\text{ff}}}$, applied to key, value, and feed-forward network (FFN) activations, respectively. Unlike LoRA, LoKr, and LoHa, which modify weight matrices, IA$^3$ directly scales internal activations. The adjusted self-attention mechanism becomes:

$$\text{Attn}(\mathbf{Q}, \mathbf{K}, \mathbf{V}) = \left( \frac{\mathbf{Q}(\mathbf{l}_k \odot \mathbf{K}^T)}{\sqrt{d_k}} \right) (\mathbf{l}_v \odot \mathbf{V}), \tag{4}$$

where $\mathbf{Q}, \mathbf{K}, \mathbf{V}$ are the query, key, and value matrices, respectively, and $d_k$ is the dimension of the key vectors. In the FFN, the adaptation is represented as:

$$\text{FFN}(\mathbf{h}') = (\mathbf{l}_{\text{ff}} \odot \theta(\mathbf{W}_1 \mathbf{h}))\mathbf{W}_2, \tag{5}$$

where $\mathbf{l}_{\text{ff}}$ is the learned scaling vector applied to the output of the first FFN layer, $\mathbf{W}_1$ and $\mathbf{W}_2$ are FFN weight matrices, and $\theta$ is the FFN activation function.

#### Fisher-transformed correlations for target-wise evaluation

Cumulative metrics on test sets provide a broad snapshot of overall model performance across all targets, but they can obscure important performance variations at the individual target level. To overcome this, we apply the Fisher $z$-transformation to both Pearson $R$ and Spearman $\rho$ correlation coefficients to enable more reliable comparisons across individual protein targets. This transformation,

$$z = \frac{1}{2} \ln \left( \frac{1 + r}{1 - r} \right), \tag{6}$$

converts the bounded correlation coefficient $r \in [-1, 1]$ into an unbounded variable $z$, which is approximately normally distributed for large sample sizes.[1] By stabilizing the variance and reducing bias inherent in raw correlation values, the Fisher transformation provides a clearer view of target-wise performance, which is critical to understanding the variability across targets in the test set.

#### Conversion between $IC_{50}$ or $K_\mathrm{d}$ and Gibbs Free Energy ($\Delta G$)

The conversion from experimental $IC_{50}$ or dissociation constant ($K_\mathrm{d}$) values to Gibbs binding free energy ($\Delta G$) was carried out using the thermodynamic relationship,

$$\Delta G = RT \ln(K_\mathrm{d} \text{ or } IC_{50}), \tag{7}$$

where $R$ is the gas constant ($R = 1.9872041 \times 10^{-3}$ kcal mol$^{-1}$ K$^{-1}$), and $T$ is the temperature in Kelvin (typically 298.15 K unless otherwise stated). When converting $IC_{50}$ to $K_\mathrm{d}$, it is assumed that $IC_{50}$ approximates $K_\mathrm{d}$ under conditions of competitive inhibition with negligible enzyme or receptor concentration compared to $IC_{50}$. $IC_{50}$ or $K_\mathrm{d}$ values provided in micromolar (µM) or nanomolar (nM) units were first converted to molar units (M) before calculating $\Delta G$.

#### Docking approach for LP-PDBBind

The LP-PDBBind[2] database provided the native receptor conformation and ligand binding poses extracted from their protein data bank (PDB) entries. When multiple alternative conformations were reported in the crystallographic data, the first alternative conformation was selected. The tautomers and protonation states of the protein-ligand complexes were obtained using the Molecular Operating Environment suite v2022-2[3] while maintaining the structural waters and cosolvents. Non-standard residues, isotopes, and cosolvents (such as crystallization buffer molecules like polyethylene glycol, ethanediol, phosphates, etc.) were appropriately removed or replaced with standard elements or residues. Coordination ions within a 6 Å radius of the ligand were retained.

For the evaluation with rDock,[4] a cavity was generated for each system using the reference ligand method. Then, the ligands underwent a single-step simplex minimization, and the `SCORE` and `SCORE.INTER` were reported. For the evaluation with AutoDock Vina v1.2.5,[5] the charges of the receptors and ligands were estimated using the Gasteiger model with MGLtools[6] and then re-scored using Vina within the automatically detected cavity.

#### References


(1) Fisher, R. Frequency Distribution of the Values of the Correlation Coefficient in Samples from an Indefinitely Large Population. *Biometrika* **1915**, *10*, 507–521.

(2) Li, J.; Guan, X.; Zhang, O.; Sun, K.; Wang, Y.; Bagni, D.; Head-Gordon, T. Leak Proof PDBBind: A Reorganized Dataset of Protein-Ligand Complexes for More Generalizable Binding Affinity Prediction. *arXiv:2308.09639* **2023**,

(3) ULC, C. C. G. Molecular Operating Environment (MOE). version v2022.2, Chemical Computing Group: Montreal, QC, Canada, 2022; `https://www.chemcomp.com`.

(4) Ruiz-Carmona, S.; Alvarez-Garcia, D.; Foloppe, N.; Gago, F.; Gervais, G.; Irwin, J.; Sverrisson, F.; Tounge, B.; Tresadern, G.; Morley, S. rDock: A Fast, Versatile and Open Source Program for Docking Ligands to Proteins and Nucleic Acids. *PLoS Comput. Biol.* **2014**, *10*, e1003571.

(5) Eberhardt, J.; Santos-Martins, D.; Tillack, A.; Forli, S. AutoDock Vina 1.2.0: New Docking Methods, Expanded Force Field, and Python Bindings. *J. Chem. Inf. Model.* **2021**, *61*, 3891–3898.

(6) Morris, G.; Huey, R.; Lindstrom, W.; Sanner, M.; Belew, R.; Goodsell, D.; Olson, A. Autodock4 and AutoDockTools4: automated docking with selective receptor flexibility. *J. Comput. Chem.* **2009**, *30*, 2785–2791.
